# Large-scale statistical mapping of T-cell receptor *β* sequences to Human Leukocyte Antigens

**DOI:** 10.1101/2024.04.01.587617

**Authors:** H. Jabran Zahid, Ruth Taniguchi, Peter Ebert, I-Ting Chow, Chris Gooley, Jinpeng Lv, Lorenzo Pisani, Mikaela Rusnak, Rebecca Elyanow, Hiroyuki Takamatsu, Wenyu Zhou, Julia Greissl, Harlan Robins, Jonathan M. Carlson

## Abstract

T-cell receptors (TCRs) interacting with peptides presented by human leukocyte antigens (HLAs) are the foundation of the adaptive immune system but population-level analysis of TCR-HLA interactions is lacking. Here we statistically associate *∼*10^6^ public TCRs to specific HLAs using the TCR*β* repertoires sampled from 4,144 HLA-genotyped subjects. The TCRs we associate are specific to unique HLA allotypes, not allelic groups, and to the paired *α*-*β* heterodimer of class II HLAs though exceptions are observed. This specificity permits highly accurate imputation of 248 class I and II HLAs from the TCR*β* repertoire. Notably, 45 HLA-DP and -DQ heterodimers lack associated TCRs because they likely arise from non-functional trans-complementation. The public class I and II HLA-associated TCRs we identify are primarily expressed on CD8^+^ and CD4^+^ memory T cells, respectively, which are responding to various common antigens. Our results recapitulate fundamental biology, provide insights into the functionality of HLAs and demonstrate the power and potential of population-level TCR repertoire sequencing.

## Introduction

The major histocompatibility complex (MHC) is a set of genes found in jawed vertebrates, which in humans encode for human leukocyte antigens (HLAs) (*1*). The primary function of HLAs is to present fragments of proteins (i.e., peptides or antigens) on the surface of cells for T cell recognition (*2*). TCRs on the surface of T cells interact with the peptides presented by HLAs (pHLA). The TCR-pHLA interaction is a key mechanism of adaptive immunity and plays a central role in the immune system’s response to infections, cancers, allergens and self-tissues targeted in autoimmunity and transplantation (*3–8*).

HLAs are both polygenic and polymorphic, allowing for a highly specific and fine-tuned adaptive immune response to diverse pathogens. The classical antigen-presenting MHC proteins— class I and class II HLAs—are found on the surface of all nucleated and professional antigen presenting cells, respectively. A large number of allelic variants have been identified across the six loci encoding class I (HLA-A, -B and -C) and class II (HLA-DP, -DQ and -DR) HLAs (*9*). Antigens presented to T cells bind to a single polymorphic *α* chain of class I HLAs (*10*, *11*) and to the *α* and *β* chain of class II HLAs (*12–14*). Both the *α* and *β* chains are polymorphic for HLA-DP and -DQ whereas only the *β* chain is polymorphic for -DR. Furthermore, the *β* chain of HLA-DR may be encoded by four loci (*DRB1*, *DRB3*, *DRB4* and *DRB5*); all individuals have *DRB1* encoded on both instances of chromosome 6 and may additionally have one of *DRB3*, *DRB4* or *DRB5* on each chromosome 6.

HLA genes can be resolved to varying degrees by sequencing and several distinct naming systems can be found in the literature. Here we adopt the 2010 WHO HLA nomenclature (*15*). For each locus, the first two fields designate the allele group and specific protein (i.e. allotype), respectively. The third and fourth fields indicate synonymous substitutions in coding and non-coding regions, respectively. For class II HLAs, the *α* and *β* chains are encoded, sequenced and typed independently. Our sequencing of HLAs lacks information on parental haplotype (i.e., phasing). In the context of molecular epidemiology, identifying (or assuming) the resolution at which causal mechanisms (and thus, clinical associations) are likely operating remains challenging. HLAs in the same allelic group tend to present similar or identical peptides (*16*), share functional properties such as relative expression levels (*17*, *18*), and serve as ligands for the same KIR receptors (*19*). Thus, many epidemiological and functional studies treat HLAs of the same allelic group interchangeably. However, structural (*20*), functional (*21*), and evolutionary evidence (*22*) suggest that, at least in some contexts, very similar HLA allotypes frequently interact with very different TCRs. A fundamental goal of this work is to establish the relationship between TCR specificity and HLA resolution and to explore the TCR specificity of class I and class II HLAs.

The human body maintains a diverse set of naive T cells where antigen specificity is determined by TCRs (*23*, *24*). These T cells are selected such that their TCRs, which are generated via V(D)J recombination, interact with pHLAs in the thymus (*25–28*). Interaction with class I and class II pHLAs directs differentiation into CD8^+^ (cytotoxic T cell) (*29*, *30*) and CD4^+^ (helper T cell) lineages (*31*), respectively. Antigen presentation by an HLA and subsequent TCR recognition in the appropriate immunological context triggers clonal expansion of naive T cells resulting in a large population of T cells expressing identical cognate TCRs (*32*). Clonal expansion of T cells with the same TCR greatly increases the chance of sampling these TCRs experimentally. As a result, subjects with matching HLAs and shared antigenic exposure have a significantly higher likelihood of sharing subsets of TCRs compared to subjects with differing HLAs and/or antigenic exposure history (*33–36*). Here we leverage this aspect of T-cell biology to identify sets of public TCRs that are over-represented in subjects sharing HLAs. We expect these sets to be enriched for HLA-restricted TCRs specific to common antigens and we use them to probe the functional nature of HLAs.

The T-cell repertoire is a rich source of information for understanding adaptive immunity (*37*, *38*). The vast majority of TCRs are heterodimers composed of an *α* and *β* chain which together encode pHLA specificity. Our data consists of T-cell repertoires of TCR*β*^1^ sequences; the paired *α* chain is unknown. While any given TCR*β* chain may randomly pair with many *α* chains, the memory T cell compartment appears to be dominated by *β* chains that pair with a single *α* chain (*39*). This observation results from the fact that TCR-pHLA binding only occurs with very specific TCR*αβ* combinations and these specific TCRs are significantly more likely to be sampled in a repertoire due to clonal expansion driven by antigen recognition. Furthermore, if the response is to a common antigen, the TCR may be observed in multiple subjects (*33*, *35*). Here we show that the public TCR*β*s we associate with specific HLAs are memory T cells targeting antigens presented by multiple subjects who share the appropriate restricting HLAs and pathogenic exposure history. Thus, these HLA associated TCR*β*s are primarily paired with compatible—albeit unknown—*α* chains for maintaining specificity to the same pHLAs and their shared specificity may be identified solely on the basis of TCR*β*.

Here we use high-throughput genetic sequencing (*40*, *41*) of the T-cell repertoires of 4, 144 subjects with HLA genotypes measured from direct-sequencing to identify *∼*10^6^ public TCRs that are statistically associated with HLA allotypes. While observing any given TCR in a repertoire may be rare, the TCR repertoire of an individual expressing a given HLA will almost always contain many TCRs that we associate with that HLA. The TCRs we associate to HLAs provide a new window into understanding the interaction between TCR and HLAs. The public nature of these TCRs and their robust HLA associations permit highly accurate imputation of class I and II HLA allotypes solely from TCR repertoires allowing us to probe functional characteristics of HLAs with respect to their TCR interactions.

## Results

### Identification of HLA-associated TCRs

Our data consist of the sequenced T-cell repertoires of 4, 144 subjects with HLAs genotyped via next generation sequencing (NGS) (*42*) (see Fig. S1 for demographic distributions). The median number of unique T cells sequenced from each individual is *∼*227, 000 and 90% of subjects have counts between *∼*74, 000 and *∼*610, 000. The majority of our samples are taken from healthy adults residing in the United States; *∼*5% and *∼*20% are Lyme and Covid positive, respectively. For a given HLA, we separate subjects into cases and controls defined as those with and without the HLA, respectively. Subjects expressing an HLA that is in the same p-group^2^ as the HLA of interest are excluded from the control group, as such HLAs have identical amino acid sequences in the peptide binding region and thus may share TCR specificity. HLA-DP and -DQ are treated as heterodimers, with cases and controls defined by *α*-*β* pairs. The *α* chain of HLA-DR is invariant and thus we treat these HLAs as monomers, similar to class I HLAs. We randomly select a fixed 80% and 20% of the samples for training and validation, respectively.

We identify sets of public HLA-associated TCRs using a statistical approach that enforces the assumption that each TCR is associated with at most one HLA allotype (we test this assumption below). This assumption enables us to disentangle the effects of linkage disequilibrium (LD) among HLA loci (*43*) that would otherwise result in a large number of spurious HLA-TCR associations (Fig. S2A). We use exact matching of the TCR*β* V-gene, J-gene and CDR3 to identify sequences that are over-represented in subjects with a given HLA allotype. Thus, our association of TCRs with HLAs is agnostic to the specific amino acid sequence, it solely relies on it being observed in multiple repertoires.

The identification procedure works as follows (see Methods for precise details): for each HLA allotype, we first create a set of candidate HLA-associated TCRs using a one-sided Fisher’s Exact Test (FET) to identify TCRs over-represented in cases. We adopt a pre-specified fiducial p-value threshold, *p^∗^*. For each unique candidate TCR, we fit an L1-regularized Logistic Regression (L1LR) model which predicts the presence of that TCR in subjects given their HLAs (represented as a binary vector of indicator variables). We tune the L1 hyperparameter *λ* to be the smallest value for which exactly one HLA parameter is non-zero. In other words, we determine which single HLA allotype best predicts the observed distribution of a given TCR in the repertoires of our training sample. We test all TCRs with p-values *< p^∗^* and retain only TCRs which associate most strongly with the HLA being modeled. As our interest here is primarily in the characteristics of *sets* of HLA-associated TCRs, we set *p^∗^* to a permissive value of *p^∗^* = 10*^−^*^4^ and use the hold-out repertoires for validation. Note that due to exclusion of p-group matched HLAs a small number of TCRs are assigned to multiple HLAs due to variations in the training data (Fig. S2B).

We associate *∼*10^6^ TCRs to specific HLAs, for a median of 2, 400 TCRs per HLA with *∼* 70% of HLAs having a total number of associated TCRs in the range of 1, 600 *−* 5, 000. To maintain consistency with other work associating TCRs with disease (*35*, *36*, *44*, *45*), we refer to these HLA-associated TCRs as *enhanced sequences* (ES).

### TCR Specificity

#### Most Enhanced Sequences are specific to HLA allotypes

To validate the HLA allotype specificity of ESs, we compare their abundance in HLA cases as compared to controls in our holdout set (see Fig. 1A,B). Overall, we find clear separation of cases and controls in the holdout data across all functional HLAs (Fig. S3-S8), highlighting the specificity of these ESs at the HLA allotype level and to the heterodimer for class II HLAs. However, there are notable exceptions.

**Fig 1:**
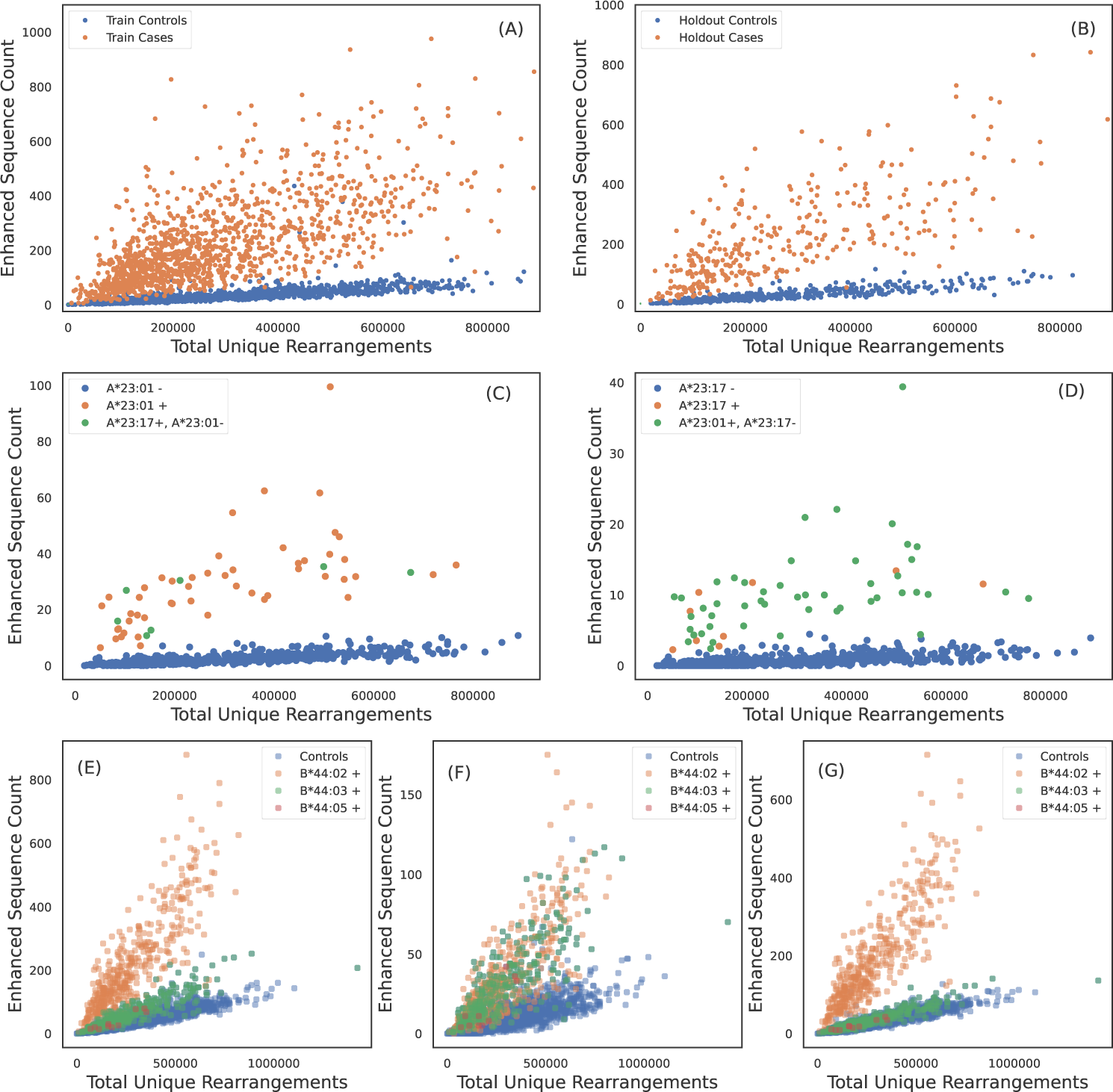
TCR specificity and apparent TCR sharing between some HLA allotypes. ESs for A*02:01 discriminate cases and controls in (A) train and (B) holdout samples. (C) A*23:01 ES counts observed in each sample as a function of sequencing depth. A*23:01 negative, A*23:17 positive subjects have counts consistent with A*23:01 positive subjects and vice-versa in (D). A*23:01 and A*23:17 appear to share the same TCR specificity. (E) B*44:02 ES counts observed in each sample as a function of sequencing depth. ESs discriminate B*44:02 positive subjects from B*44:02 negative subjects. However, B*44:03 and B*44:05 positive subjects (green dots) have elevated counts as compared to controls (blue dots). (F) ES counts plotted against sequencing depth for the subset of ESs which associate more strongly with the B*44 group as compared to B*44:02 allotype (identified using the L1LR association method). The subset of ESs in (F) elevate all B*44 positive subjects equally suggesting the TCRs in this ES subset are specific to the three allotypes in the group. (G) ES counts plotted against sequencing depth for the subset of ESs which associate only to B*44:02, i.e., this set excludes the ESs shown in (F). ESs plotted in (G) clearly separate B*44:02 positive subjects from B*44:02 negative subjects, including other allotypes in the B*44 group.

We identify one HLA class I allotype pair (Fig. 1A-1B) and 11 class II pairs (Table 1) that appear to completely share TCRs despite differing in their two-field designation. We refer to these as “degenerate” HLAs. For each of these degenerate pairs, ESs specific to one HLA are equally distributed among individuals expressing either HLA (Fig. 1C-D). Ten of the twelve degenerate HLAs we identify have amino acid differences in a single position not in the peptide presentation and TCR binding domain and thus are in the same p-group.

To further validate our assumption that most of these TCRs are specific to HLA allotypes, not allelic groups (with the exception of noted degenerate pairs), we use the L1LR method to assign TCRs to either the allelic group (1-field) or the allotype (2-field). We restrict our analysis to class I HLA groups observed in *>* 200 subjects with the most common allotype representing *<* 70% of subjects. Among the six allelic groups tested, we find five have a negligible fraction (*∼*1%) of ES associated to the group (A*30, A*33, A*68, B*15, B*35, C*07), indicating that the majority of HLA-associated TCRs we identify are allotype specific.

We find one HLA group where a subset of TCRs in the ES set appear to be specific to multiple allotypes in the group: B*44. We find that 10% of TCRs originally identified as B*44:03-specific are assigned to the B*44 group (Fig. 1E-1G; similar conclusions are reached when starting with the less prevalent B*44 allotypes). This set of B*44-specific TCRs segregate all B*44 positive from negative individuals in the holdout (Fig. 1F), while the remaining *∼*90% of TCRs originally identified as B*44:03-specific separate B*44:03-expressing individuals from those who express B*44:02 or 44:05 (Fig. 1G). These three B*44 allotypes differ in only two amino acids (residue 140 and/or 180), and B*44:02 and 44:03 are known to share a large fraction of their peptide repertoire and some of their TCR repertoire (*46*).

Degenerate HLAs that completely share their TCR repertoire typically differ in one amino acid outside the binding domain. Similarly, B*44 has two polymorphic positions outside the binding domain and displays a high degree of sharing. On the other hand, groups in which we observe no TCR sharing tend to have one or more amino acid differences in the binding domain. Thus, the degree of sharing we observe appears to be correlated to the number of differing residues and the position at which the differences occur. Taken together, these results show that many TCRs (indeed, the vast majority of those identified here) are specific to distinct HLA allotypes, regardless of shared peptide repertoire or binding domain similarity. Thus, many characteristics of TCR-pHLA interactions differ among highly related HLA allotypes. As expected, no such specificity was observed among HLA allotypes that differ only in synonymous substitutions (ie, at 3- and 4- digit resolution; Fig. S9). We note that to mitigate the effects of linkage disequilibrium, our algorithm for identifying ESs assumes TCRs are specific to individual HLA allotypes (see methods). Thus, our methodology is not designed to identify an unbiased set of TCRs specific to multiple HLA allotypes. Our identification of a small fraction of ESs with specificity to multiple HLA allotypes warrants a systematic investigation which is beyond the scope of this work.

#### Most TCRs are specific to class II heterodimers, not subunits

Functional class II HLAs are stable heterodimers with both the *α* and *β* chain contacting the peptide. As such, we expect class II HLA-associated TCRs to be specific to the heterodimer and not the protein subunits (i.e. the *α* or *β* chains individually). To directly test this hypothesis, we use the L1LR method to determine if a TCR is more strongly enriched among individuals expressing both the *α* and *β* chains or individuals expressing only one or the other subunit. Across 37 heterodimers, we find that *∼*146000 (70%), *∼*20500 (10%) and *∼*43000 (20%) of ESs are most strongly associated with the heterodimer, alpha and beta subunits, respectively. We note that not all 37 heterodimers exhibit single-chain specificity (see below). Thus, the vast majority of class II associated TCRs appear to be specific to the combined *α − β* chains. This finding is bolstered by the fact that the ES sets we derive discriminate HLA-DP and - DQ heterodimers and not individual subunits (Fig. 2; see also Fig. S6-7, which show ES distributions for all heterodimers).

**Fig 2:**
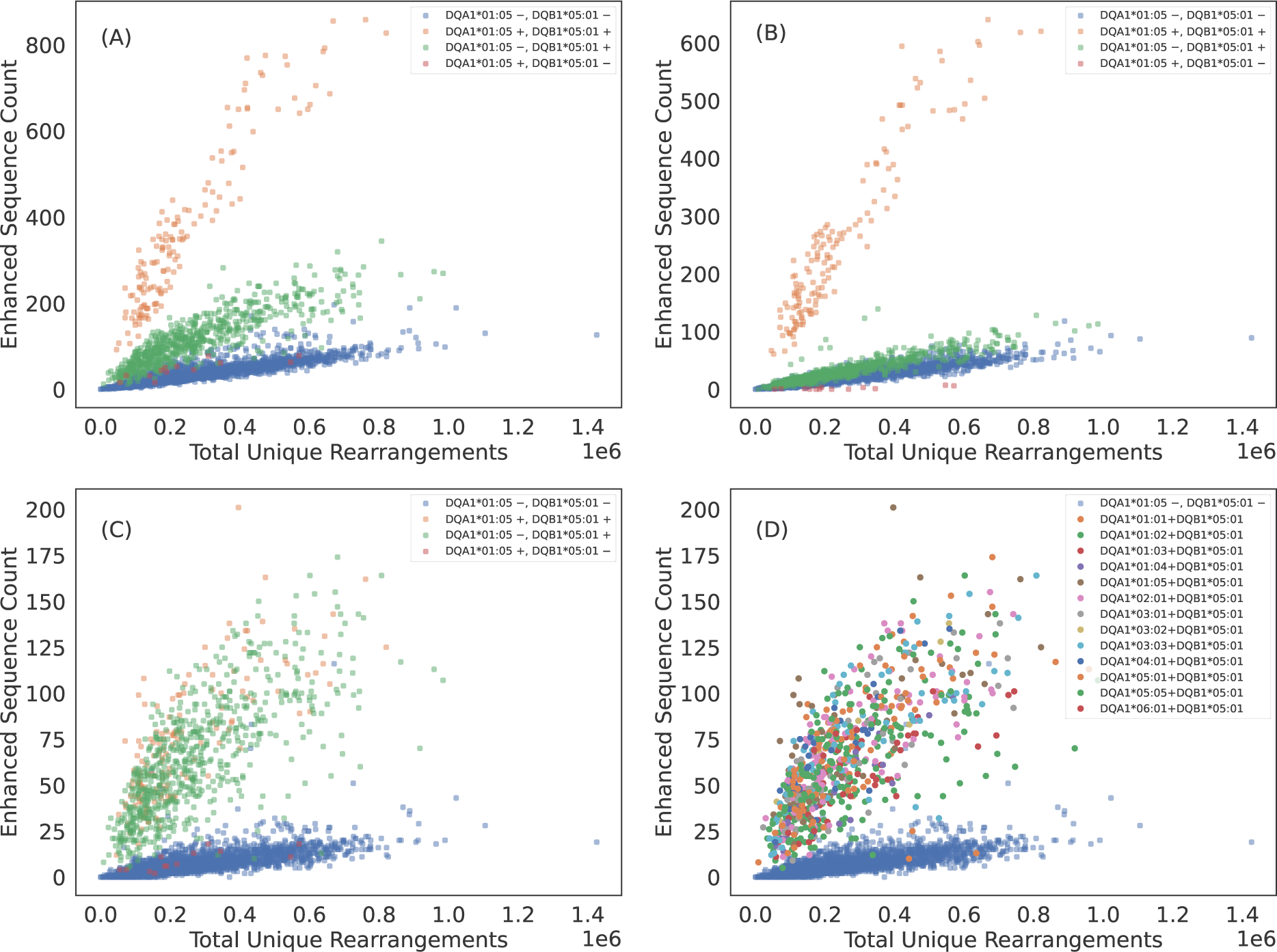
Some TCRs appear to be specific to the class II HLA subunit rather than heterodimer. (A) DQA1*01:05+DQB1*05:01 ES counts observed in each sample as a function of sequencing depth. The ESs separate DQA1*01:05+DQB1*05:01 positive subjects from DQA1*01:05+DQB1*05:01 negative subjects. However, subjects who have the DQB subunit appear elevated above the control population of subjects with neither subunit. (B) ES count plotted against sequencing depth for the subset of ESs which associate most strongly with DQA1*01:05+DQB1*05:01 heterodimer via the L1LR method. A majority of TCRs which make up the ES set for DQA1*01:05+DQB1*05:01 appear to be specific to the heterodimer. (C) ES count plotted against sequencing depth for the subset of ESs which associate most strongly to subunit DQB1*05:01 via the L1LR method. A small fraction of TCRs which make up the ES set for DQA1*01:05+DQB1*05:01 appear to be specific only to the *β* chain subunit. (D) Same as (C) but color-coding subjects with DQB1*05:01 by their various *α* chain pairings. This subset of TCRs appear to be specific to all possible heterodimeric combinations which include DQB1*05:01, thus suggesting specificity solely to the *β* chain subunit. Results are shown for train sample and are consistent with holdout sample.

We find that only a subset of ESs and HLAs exhibit exceptions to heterodimeric specificity. For example, some TCRs appear to be associated to all heterodimers composed of the DQB1*05:01 subunit, indicating many of these TCRs are associated with the subunit itself (Fig. 2). DQB1*05:01 is the clearest example of TCR specificity to a subunit. However, we observe such subunit specificity across multiple subunits: DPB1*01:01, DQB1*02:01 (DQB1*02:02)^3^, DQB1*03:01, DQB1*05:01, DQB1*06:03, DQA1*03:01 (DQA1*03:03) and DQA1*05:01 (DQA1*05:05). TCR specificity to subunits appears to be more common for the HLA *β* chain and to the HLA-DQ locus, though we observe single-chain specificity in both the *α* and *β* chains of HLA-DQ and a *β* chain of HLA-DP. We note that our identification is likely not exhaustive as many heterodimers lack enough diversity in one or both subunits to statistically associate TCRs independent of the heterodimer.

Based on a limited set of solved structures, a conserved binding pattern has been proposed for class II TCR-pHLA interactions such that TCR*α* contacts the *α* helix of the HLA *β* chain and TCR*β* contacts the *α* helix of the HLA *α* chain (*47*). Thus, the various specificity patterns we observe may reflect TCR interactions strongly mediated by peptides and not the direct interactions between TCRs and HLAs which may be conserved. Future analyses based on larger sets of solved or perhaps *in silico* generated structures of class II TCR-pHLA complexes will be informative for exploring the structural basis of the various specificity patterns we identify.

### TCR breadth is proportional to zygosity

HLA homozygosity has been epidemiologically linked to poor clinical prognosis in the context of both chronic HIV infection (*48*) and cancer checkpoint-inhibitor immunotherapy (*49*), possibly due to the reduced size of the HLA-restricted peptide repertoire available for T-cell recognition. This reduced size of the antigen repertoire, coupled with higher relative surface concentration of pHLAs associated with the homozygous protein, may directly impact the TCR repertoire by increasing the probability of clonal expansion of T cells expressing cognate TCRs. Consistent with this hypothesis, we find that the distribution of ES counts is elevated among homozygous individuals (Fig. 3A-3F). Across all HLAs, the distribution of ESs is (on average) about one standard deviation higher for homozygous as compared to heterozygous individuals (Fig. 3G,H). Notably, the ES distribution is an additional standard deviation higher among individuals homozygous at both the *α* and *β* locus for HLA-DP or -DQ (“double homozygous”, Fig. 3H).

**Fig 3:**
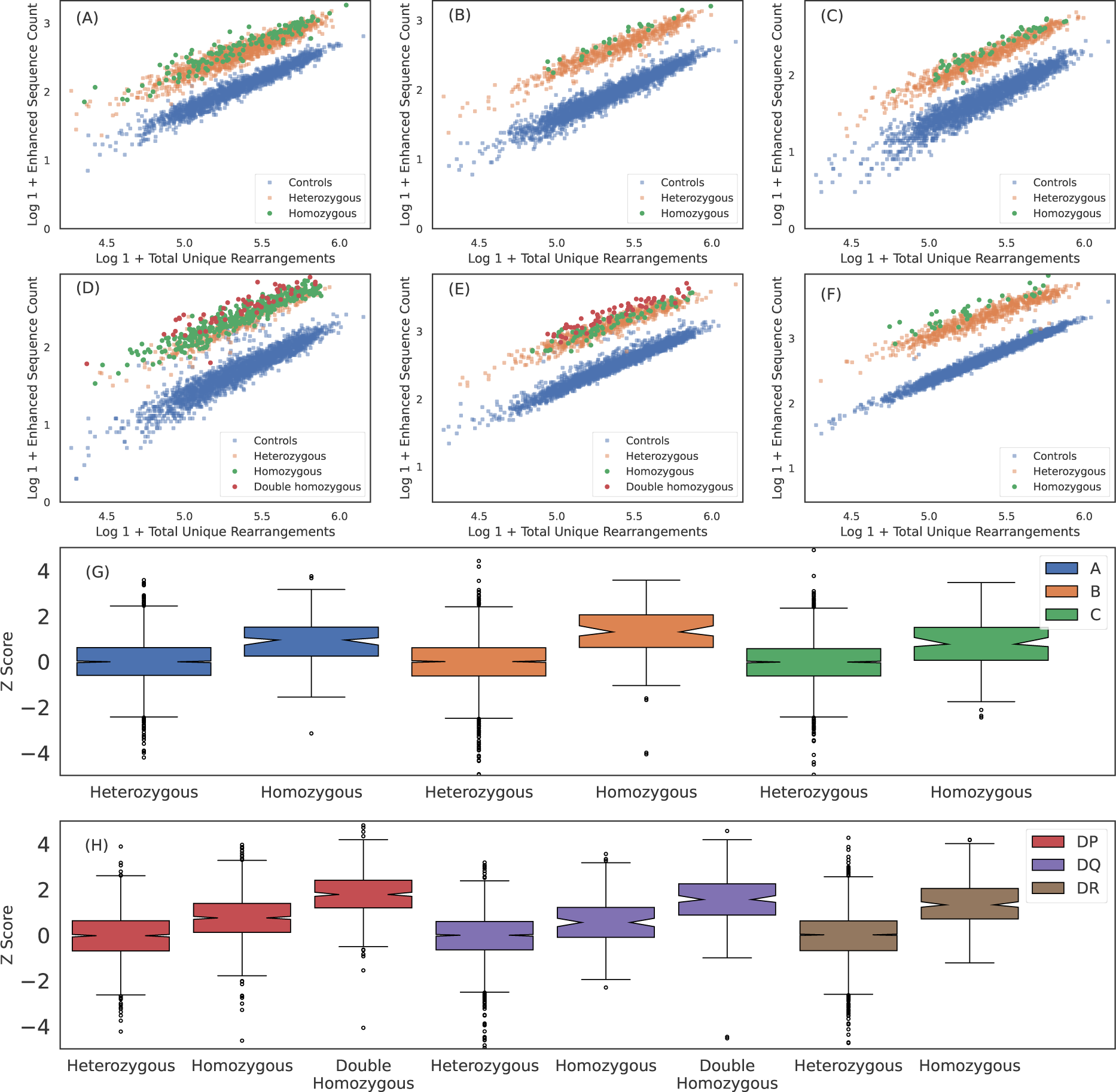
Breadth of T cell response is proportional to HLA zygosity. ES counts for (A) A*02:01, (B) B*07:02, (C) C*04:01, (D) DPA1*01:03+DPB1*02:01, (E) DQA1*01:02+DQB1*06:02 and (F) DRB1*07:01 ESs observed in each sample plotted against total number of unique rearrangements. HLA negative, heterozygous positive and homozygous positive subjects are shown in blue, orange and green, respectively. For HLA-DP and DQ, double homozygous subjects are shown in red. The breadth of the ES response appears to be correlated with homozygosity across all six loci. We quantify the the increased breadth resulting from homozygosity by fitting the mean and standard deviation of the ES counts in heterozygous cases for each HLA as a function of sequencing depth and then calculating the z-score for all subjects and all well-represented HLAs. We aggregate z-score distributions per-loci for (G) class I and (H) class II HLAs. Results are shown for train sample and are consistent with holdout sample.

While these results are consistent with increased relative antigen abundance increasing clonal expansion of associated T cells, an alternative hypothesis is that a decrease in the relative surface expression or decreased antigenic diversity in homozygous subjects results in less crowding out by other HLAs or TCRs, respectivly. This crowding hypothesis implies an increased breadth across multiple loci within a given class for homozygous subjects. We test this hypothesis by examining whether a homozygous subject at one locus has higher breadth at another class-matched loci. For example, if HLA and/or TCRs crowd each other, we expect that a subject homozygous for HLA-A will also have a higher average breadth in their HLA-B and/or HLA-C response due to lower diversity at the class I loci. We do not find evidence of such an effect (see Fig. S10) and thus conclude that crowding out by HLAs and/or TCRs may not be the primary driver of increased breadth unless it is restricted to a particular locus.

Taken together, these results suggest homozygosity at a particular locus increases the breadth of the T-cell response against peptides presented by that HLA, possibly through increased surface expression and antigen presentation.

### Imputing HLA genotype from TCR repertoires

The clear separation of ES counts in HLA cases versus controls implies that HLA allotypes can be easily imputed from HLA-associated TCRs alone. To this end, we fit a simple logistic regression model for each HLA allotype observed in at least 30 training samples (representing *∼*1% expression frequency), predicting whether an individual expresses that HLA as a function of observed ES and total-unique-rearrangement (log) counts (see Fig. 4A-4B for representative examples, Figs. S3-S8 for all HLAs tested).

**Fig 4:**
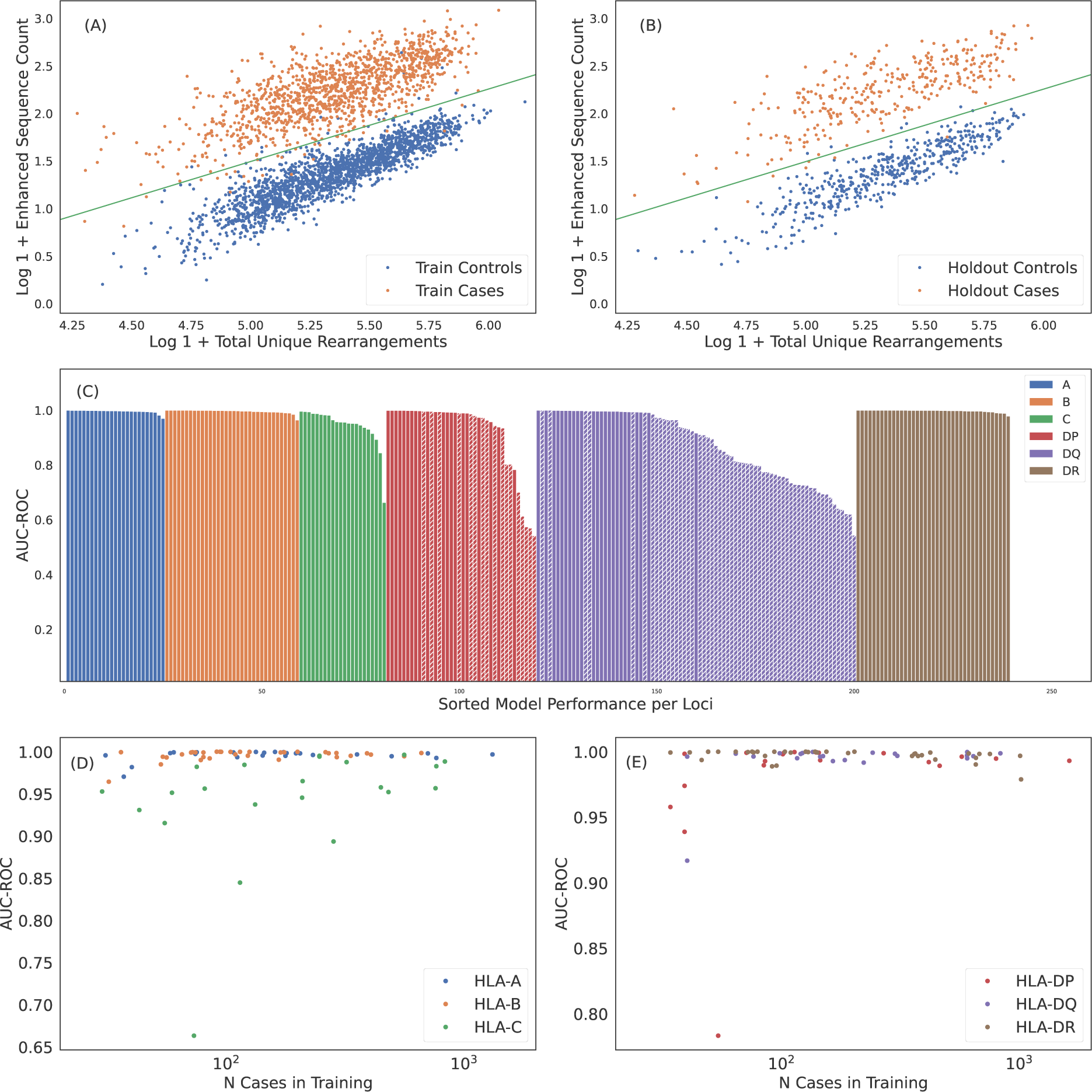
Robust predictions of hundreds of HLAs solely based on public T cells observed in the repertoire. ES counts for A*02:01 cases and controls plotted against total number of unique rearrangements for (A) train and (B) holdout samples. The green line indicates the call threshold. (C) AUC-ROC of HLA models across each loci sorted by performance. The hashed bars for HLA-DP and -DQ indicate heterodimer resulting from subunits combinations in linkage equilibrium suggesting trans-complementation. We show below that many of these HLAs are likely non-functional. Performance of (D) class I and (E) class II HLA models as a function of the number of training samples color-coded by loci. Performance correlates with the number of training samples, as expected. HLAs shown by hashed bars in (C) are excluded in (E). In (C)-(E) we show performance derived from 5-fold cross validation (CV) of the train sample. CV performance is consistent with holdout performance.

Over all the imputation accuracy is extremely high with area under the receiver operating characteristic curve (AUC-ROC) scores of *≥* 0.9 for all but 3 of the 120 HLA-A,-B, and -DR allotypes modeled. This accuracy highlights the specificity of HLA ESs, indicating HLAs at these loci can be accurately imputed from immunosequencing alone. Furthermore, this accuracy confirms that HLA*β* alone is typically sufficient to identify shared antigen specificity of public T cells. Among class I HLAs, HLA-C allotypes are a notable outlier with relatively lower classification performance even among models with a significant number of positive training examples (Fig. 4D). We hypothesize that this reduced performance is due in part to the *∼*10*×*-lower surface expression of HLA-C compared to allotypes expressed by other class I genes (*50*).

Model performance among HLA-DP and -DQ heterodimers is substantially more variable than performance at the other loci. We find that 8 of the 30 HLA-DP and 37 of 81 of HLA-DQ fail to achieve AUC-ROC scores of *>* 0.9 even among heterodimers expressed in a high number of individuals in our training population (Fig. 4E). Notably, HLA-DQ and -DP are the only loci we study with polymorphic *α* chains, suggesting that heterodimer incompatibility may explain the lack of associated TCRs and the corresponding inability to impute expression of these heterodimers. We explore this issue in the next section.

We confirm the generalizability of HLA imputation across genetic and geographic (and thus, possible antigenic) backgrounds by assessing accuracy of HLAs imputed by our model compared to sequence-based HLA typing among an independent cohort of 136 individuals from Kanazawa Japan. For this analysis we use a set of 135 HLAs across all six loci that yield models with cross-validation precision *>* 0.9, recall *>* 0.8 and *≥* 30 positive training cases. Applying this set of models to the Japanese cohort results in an average of 8.5 imputations per individual as compared to 11.0 per individual in our holdout cohort. This difference in the number of imputed HLAs reflects the shift in genetic background of the Japanese cohort as compared to the training data (see Fig. S1 for self-reported ethnicity in training data). Nevertheless, we observe high model accuracy, with an overall F1 score^4^ aggregated across all imputations in the Japanese cohort that is comparable to that observed in the original holdout cohort (0.925*±*0.006 and 0.940 *±* 0.002^5^, respectively).

### Poor-performing class II models are trans-complemented, non-functional HLAs

HLAs are inherited as a haplotype, such that one full set is inherited on a single chromosome from each parent (*51*). The *α* and *β* chains of HLA-DP and -DQ^6^ pair after synthesis, yielding a phenotype of up to four unique heterodimers in each individual: two formed in *cis*, where both subunits are encoded on the same chromosome, and two in *trans*, where the subunits are encoded on opposite chromosomes. Given the high degree of polymorphism observed in HLA-DP and -DQ subunits, it is perhaps unsurprising that some pairs of *α* and *β* chains do not form stable heterodimers (*52–54*). Based on structural and sequence analysis of HLA-DQ, Tollefsen et al. (*54*) propose specific group pairings that likely form stable heterodimers (though they note a small number of exceptions to the pairing rules).

In the context of the present study, the proposed existence of incompatible (and thus non-functional) *α* and *β* chains implies two testable hypotheses: (1) that co-inheritance of incompatible pairs on the same chromosome will be under strong negative selection (*54*); and (2) that incompatible pairs are unable to elicit a T-cell response and thus will not be associated with any public TCRs.

The first hypothesis implies that co-expression of incompatible pairs almost always results from *trans*-complementation; as such, incompatible pairs will be in linkage equilibrium since they are not co-inherited. Conversely, *cis*-complementation should necessarily result in functional pairs thus all pairs forming from subunits in linkage disequilibrium should be functional. For two subunits *α* and *β*, with respective expression frequencies *f_α_*and *f_β_*and co-expression frequency *f_α_*_+_*_β_*, linkage equilibrium results in an expected co-expression frequency of approximately *E_LE_*[*f_α_*_+_*_β_*] = 2*f_α_f_β_* (*55*). Overall, we find that a majority (75 of 119) HLA-DP and -DQ *α*+*β* pairs appear to be in linkage equilibrium, i.e., not co-inherited (Fig. 5A; only subunits comprising heterodimers observed in at least 30 individuals are considered). Notably, all of the pairs that Tollefsen et al. (*54*) propose to be incompatible are in equilibrium (Fig. 5B).

**Fig 5:**
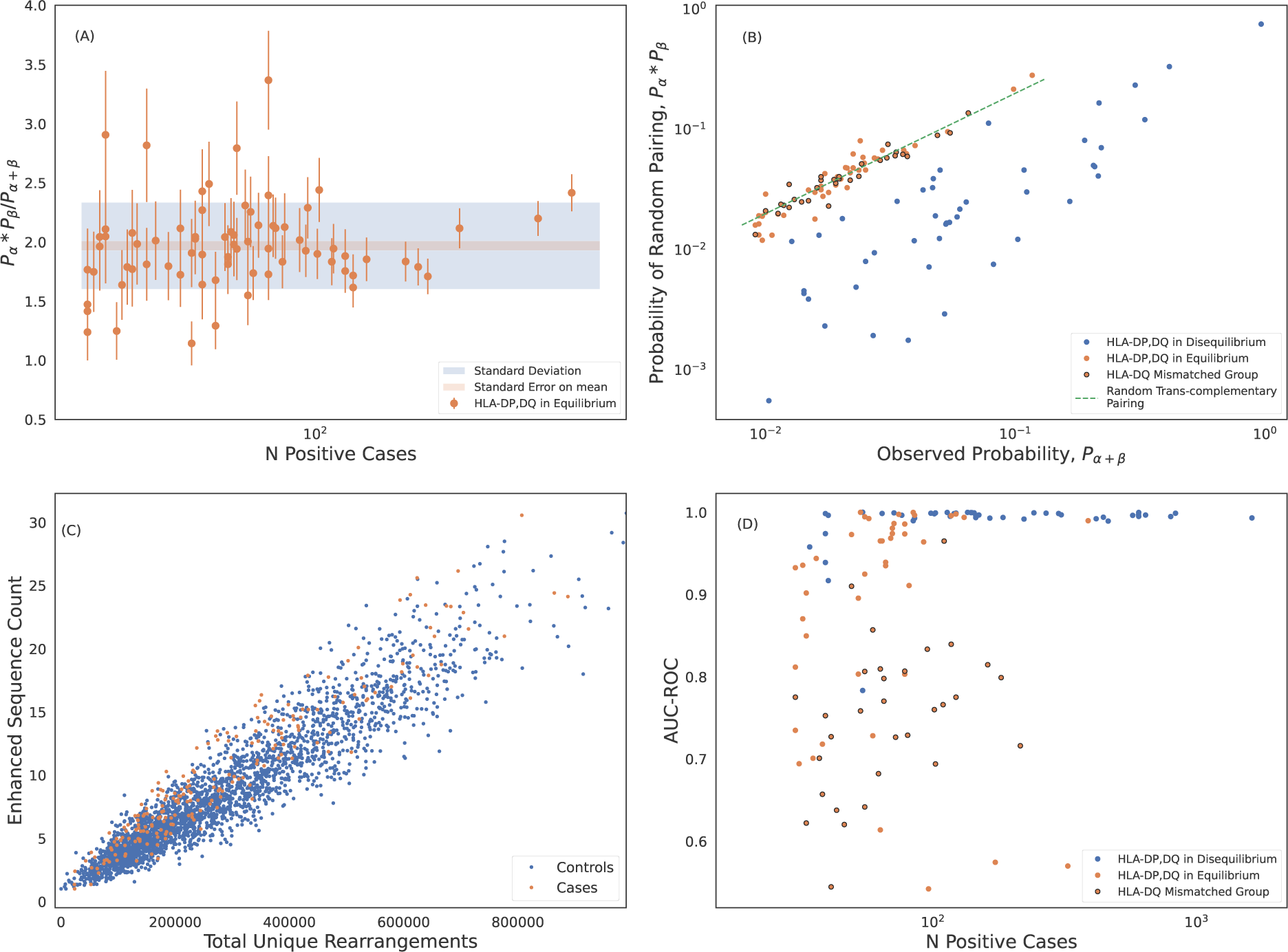
Trans-complemented heterodimers may not form stable HLAs. (A) Statistical analysis of the marginal-to-joint probability ratio of heterodimers forming from subunits in linkage equilibrium. We identify these heterodimers as those which differ *>* 5*σ* from ratio of 2 expected for genes in linkage equilibrium. We measure an average probability ratio of 1.97 *±* 0.04. Error bars are derived from propagating poisson uncertainties. (B) Expected probability of randomly pairing *α* and *β* chains of DP and DQ heterodimers plotted against the observed joint probabilities. Probabilities are calculated from the normalized inverse frequencies. The probability of randomly pairing is calculated as the product of the observed marginal probabilities of the subunits. The dashed green line is the expected correlation for random trans-complementary pairing. We identify hetorodimers formed from subunits in linkage equilibrium (shown in orange) as in (A). DQ heterodimers formed from pairing of mismatched groups as defined by (*54*) are shown with black circles. These mismatched group heterodimers cluster around the dashed green line indicating random trans-complementary pairing. (C) DQA1*01:02+DQB1*03:01 is an example of an HLA where we are unable to identify any ESs that separate cases and controls. (D) Model performance of HLA-DP and -DQ models including HLAs forming from subunits in linkage equilibrium. Here we show performance derived from 5-fold cross validation (CV) of the train sample which is consistent with holdout performance.

Given the tight linkage between genes encoding subunits, any pair expressed in *cis* in at least one individual is likely to be in LD in a large cohort. Consistent with this hypothesis, every individual heterozygous at both the *α* and *β* locus has at least two of four *α/β* pairs in LD; conversely, no individual should have more than two *α/β* pairs in equilibrium. We confirm that no subject in our cohort has more than two heterodimers formed from subunits in linkage equilibrium, as expected. Taken together, we conclude that LD is a strong proxy for *cis*-complementarity, and that, given strong selection pressure, incompatible pairs are only expressed in *trans*.

If incompatible pairs are truly non-functional (with respect to antigen presentation and T-cell recognition), then there should not be any TCRs that are specific to such pairs. As an example, we are unable to identify distinguishing TCRs for DQA1*01:02+DQB1*03:01, which both violates the Tollefsen et al. (*54*) pairing rules and is in linkage equilibrium (Fig. 5C). Moreover, all of our high-frequency, poor-performing HLA-DP and -DQ imputation models are for heterodimers forming from subunits that are in linkage equilibrium, and thus are almost certainly expressed in *trans* (Fig. 5D; see also Fig. 3C). Moreover, almost all pairs that violate the Tollefsen et al. (*54*) pairing rules have lower-than-expected imputation performance (Fig. 5D).

Notably, while heterodimer incompatibility implies both linkage equilibrium and poor model performance, not all heterodimers forming from subunits in linkage equilibrium result in poor performing models. Thus, trans-complementation may yield heterodimers which can drive a T-cell response resulting in identifiable TCRs and a high-accuracy imputation model (Fig. 5D).

Using these results on model performance and gene linkage, we can extend the Tollefsen et al. (*54*) pairing rules: DPA1*02 appears to form unstable heterodimers when paired with DPB1*02 and DPB1*04; DPA1*01 is apparently unrestricted, forming stable heterodimers with subunits from all DPB1 groups in our sample.

### HLA associated sequences are memory T cells responding to common antigens

The T-cell repertoire consists of a mixture of naive and memory T cells. In principle, HLA-restricted antigen presentation will bias both thymic selection and clonal expansion, and thus HLA-specific signatures may exist in both compartments. However, our statistical approach to identifying HLA-specific public TCRs likely favors identification of TCRs from memory T-cells.

We investigate characteristics of HLA associated sequences by comparing the distribution of ESs observed in the memory and naive compartments sequenced from 45 individuals who were not included in the original study. For each subject, we sequence five separate repertoires: the memory and naive compartments of CD8^+^ and CD4^+^ T cells, respectively, and the unsorted repertoire. As sequence-based typing was unavailable for these individuals, we treat the imputed HLAs from the the unsorted repertoire as ground truth (limiting to 90 models with *>* 0.95 precision and recall). For each repertoire and each possible HLA, we compute the weighted breadth of the HLA-specific ESs observed in the repertoire. Across all 45 unsorted repertoires, the weighted breadth of both class I and class II ESs is substantially higher for the HLAs an individual expresses compared to those they do not express (Fig. 6A; compare HLA-positive to -negative).

**Fig 6:**
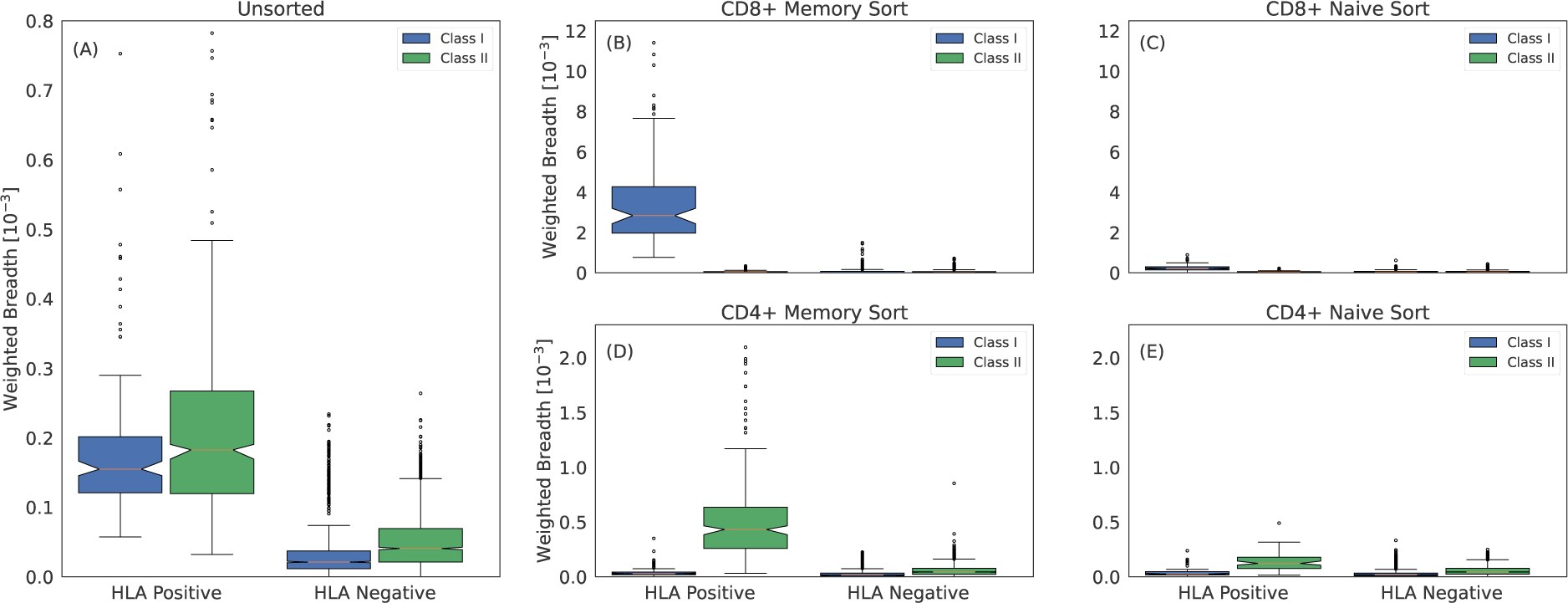
Class I and class II associated ESs are TCRs from CD8^+^ and CD4^+^ memory T cells, respectively. (A) Weighted breadth of 90 HLA associated ESs measured in 45 subjects with imputed HLAs. The breadth is sorted by whether the subject has the HLA or not and is then aggregated across all subjects and HLAs. To generate a comparable breadth across HLAs, which have varying number of ESs, we measure the median breadth in controls for all 90 HLAs, which we rescale to a mean of 1. We then normalize the breadth of any given HLA by this value. Breadth measured in (B) CD8^+^ memory sorted, (C) CD8^+^ naive sorted, (D) CD4^+^ memory sorted and (E) CD4^+^ naive sorted repertoires. We measure slightly elevated breadth in the CD8^+^ and CD4^+^ naive sorted repertoires for class I and class II HLAs, respectively. This elevation may be due to surface markers not perfectly discriminating naive and memory compartments or to a weak HLA specific signal due to the HLA interactions required for maintaining homeostatic equilibrium of naive T cells (*65*).

Within the sorted compartments, a striking pattern emerges: in the naive compartments, there is little difference in the breadth of ES specific for an individual’s expressed HLAs compared to background (Fig. 6C,E), while in the memory compartments, ESs specific to the individuals’ expressed HLAs have far higher breadth (Fig. 6B,D). Moreover, within the memory compartment, CD8^+^ cells are closely linked to high relative breadth of class I HLA ESs, while CD4^+^ cells are closely linked to high relative breadth of class II HLA ESs. This result provides further confirmation that ESs are correctly mapped to HLAs despite the challenges of HLA LD.

The centrality of the memory compartment in driving our HLA imputation signal raises several additional hypotheses. The first is that clonal expansion increases the likelihood of detection for ESs. This increased likelihood implies that, while ESs are frequently observed in individuals without the associated HLA, they will tend to be at higher repertoire frequency among individuals who do express the HLA. Indeed, we observe a notable increase in the distribution of clonal frequency among cases compared controls (Fig. 7A).

**Fig 7:**
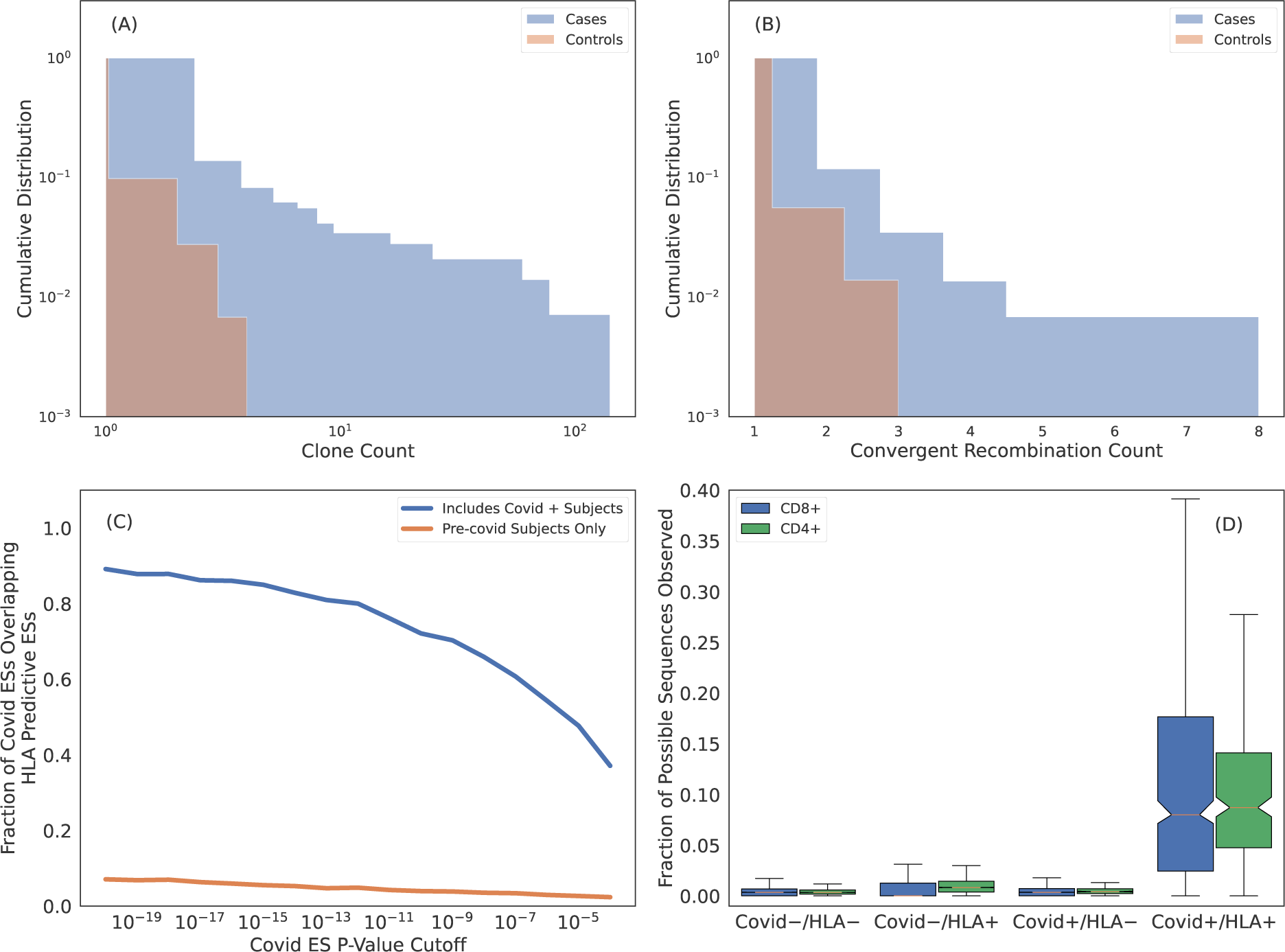
Many HLA ESs are T cells responding to common pathogens. (A) Cumulative distribution of clone count per unique rearrangement as measured in cases (blue) and controls (orange). ESs are generally more expanded when they are observed in subjects with the restricting HLA. (B) Cumulative distribution of the number of unique rearrangements mapping to an ES, i.e., convergent recombination count. ESs show more convergent recombination when observed in cases (blue) as compared to controls (orange). (C) Intersection of SARS-CoV-2 specific ESs derived via a FET on samples with PCR labels and HLA ES sets. Blue curves are HLA ES sets derived from samples which include Covid-19 positive subjects and orange curves are from samples with all Covid-19 positive subjects removed. (D) Fraction of SARS-CoV-2 ESs observed in subjects relative to the total fraction possible given the ES HLA association determined from intersecting the SARS-CoV-2 ESs with the HLA ES sets. Here we impute HLAs using our models. For “HLA +” we only count sequences associated with HLAs inferred for the subject and for “HLA -” we only count sequences associated with HLAs that are not inferred for the subject.

The second hypothesis is that clonal expansion results from antigen exposure, which will also result in polyclonal expansion of T cells that express distinct TCRs that all respond to the same antigen. An extreme form of polyclonal expansion is *convergent recombination*, in which multiple, distinct TCR DNA sequences encode identical amino acid sequences. Consistent with this hypothesis, the per-ES convergent recombination rates are higher when ESs are observed in cases as compared to controls (Fig. 7B; conditional on the ES being present in the repertoire).

The publicity of HLA ESs combined with their apparent antigen-specific clonal expansion suggests that these sequences are responding to antigens which are common in the human population (*56*). We test this hypothesis in the context of SARS-CoV-2 exposure. Because of its novel nature, SARS-CoV-2 is especially well-suited for this task as we are able to confidently assign Covid-19 negativity to samples collected before 2020. In our training sample for deriving HLA restricted sequences, 694 of the 4, 144 subjects (*∼* 20%) are Covid-19 positive based on PCR labels; the remaining samples are collected before 2020 and thus Covid-19 negative. Because SARS-CoV-2 exposure is relatively high in our training sample, we expect that some of the HLA-specific ESs we identity are SARS-CoV-2 specific.

We identify 6866 SARS-CoV-2-specific ESs using a set of 1523 SARS-CoV-2 PCR-positive samples that have no overlap with the typed HLA training samples and 4386 controls (1008 controls overlap with the typed HLA samples). A high proportion (80% of the most confident SARS-CoV-2 specific ESs) are also HLA-specific ESs (Fig. 7C, blue line). This overlap in Ess results from the fact that *∼* 20% of our typed HLA training samples are SARS-CoV-2 positive, i.e., SARS-CoV-2 specific ESs are predicting of HLAs as well. In contrast, if we define an alternative set of HLA-ESs using only 3, 450 repertoires which were sampled prior to 2020, we find very little overlap (Fig. 7C, orange line). The limited overlap we do observe may be due to cross-reactivity to homologous epitopes from other coronaviruses and/or false positive sequences in the SARS-CoV-2/HLA ES set.

The intersection of SARS-CoV-2 specific ES set with the and HLA specific ES sets yields 1, 880 TCRs (using a threshold of *p^∗^* = *p <* 10*^−^*^4^) that are associated with a particular HLA in the context of SARS-CoV-2 infection. Thus, these ESs are almost certainly specific to commonly targeted SARS-CoV-2 T-cell epitopes. To confirm this specificity, we impute HLAs for samples in the SARS-CoV-2 training data^7^ and then compute the fraction of HLA-associated SARS-CoV-2 ES observed in their repertoire. For “HLA +” and “HLA -” subjects we only count sequences if the subject has or does not have the HLA which the SARS-CoV-2 ES is associated with, respectively. Notably, both class-I and class-II associated TCRs are far more commonly observed in individuals with the restricting HLA and known SARS-CoV-2 infection (Fig. 7D), thus confirming the HLA- and pathogen-specificity of these TCRs and further demonstrating that while TCRs may be cross-reactive to many distinct antigens, the likelihood of cross-reactivity *in vivo* is low (*57*).

We conclude that, taken together, these results demonstrate that the majority of HLA-specific ESs are the result of clonal expansion of T cells responding to common antigens, thus establishing an immunological foundation for HLA imputation from TCR repertoires.

## Discussion

The T-cell repertoire of any individual consists of *∼*10^8^ unique TCRs out of an estimated *∼*10^16^ possibilities (*58*, *59*), of which only 10^5^ *−* 10^6^ TCRs may be practically sampled from any single repertoire using current techniques. Given the enormous diversity of possible TCRs, naively, little overlap in the TCR repertoire of different subjects may be expected. However, several factors significantly increase the likelihood of observing public TCRs: (1) the probability distribution of TCRs generated via V(D)J recombination is non-uniform and spans *∼*25 orders of magnitude (*60*), such that higher generation probability TCRs are more commonly observed in the naive repertoire; (2) antigen-experienced T cells clonally expand and are thus more likely to be observed in a repertoire; and (3) the immune response is focused on only a few immunogenic epitopes per HLA out of the many possible derived from any given antigenic exposure, an effect called *immunodominance* (*61*). As a consequence, the likelihood of observing the same TCRs in the repertoires of multiple subjects with shared antigenic exposure and appropriate restricting HLA is significantly higher than naively expected. Studies have shown that public TCRs can be identified that permit sensitive and specific diagnosis of individuals with past SARS-CoV-2 infection (*35*) and Lyme disease (*36*) as well as to determine who is seropositive for Cytomegalovirus (*33*). Given that these antigens are presented by HLAs, it is perhaps unsurprising that these public TCRs are also specific to the HLA context (*33*, *34*). Here we leverage this public fingerprint of TCRs to identify HLA associated sequences, allowing us to impute HLA types with extremely high accuracy and opening a new window into functional characteristics of HLAs.

We identify HLA specific TCRs using only the TCR*β* repertoire, even though the specificity of pHLA is to TCR*αβ*. Enhanced sequences elicit an immune response only in subjects with the appropriate shared restricting HLA and pathogenic exposure history (Fig. 7D), meaning, enhanced sequences are responding to the same pHLAs in different subjects. This result implies that an observed TCR*β* shared among multiple subjects in a given HLA context is very likely paired with one or a very small set of compatible TCR*α*s. In this respect, TCR*β* repertoires are not unique; analysis of TCR*α* repertoires would lead to similar results and conclusions as presented here^8^. Thus, our ability to identify HLA specific TCRs using only a single chain of the TCR heterodimer is not a characteristic of the TCR*β*s but rather a consequence of T cell immunology and our procedure for identifying enhanced sequences.

A key finding of this work is that TCRs are typically specific to HLA allotypes (two field resolution) and to class II heterodimers encoded by both the *α* and *β* chains, although there are some notable exceptions of TCRs that are specific to HLA groups (one field resolution) or to either the *α* or *β* subunits of HLA-DP and -DQ. While most HLAs elicit a strong and diverse public TCR response, others elicit little or no response, consistent with these HLAs deriving from incompatible *α* and *β* subunits and providing further support for the widespread prevalence of non-functional class II heterodimeric pairs resulting from trans-complementation. Furthermore, we find that the breadth of an HLA-specific TCR response is larger among individuals expressing two (or more) copies of the HLA, suggesting a dose-dependent effect of antigenic exposure on the diversity of expanded T-cell clones. These insights highlight the exquisite specificity of public TCRs and demonstrate the potential of population-level TCR analysis for probing functional aspects of the immune system.

We show that class I and class II HLA-associated TCRs are found on CD8^+^ and CD4^+^ memory T cells, respectively, which is consistent with their publicity resulting from clonal expansion in response to antigenic-exposure. The public nature of these TCRs suggests that they are likely specific to peptides derived from common pathogens, vaccines and conserved endogenous-antigens. Consistent with this hypothesis, *∼*20% of subjects in our training sample are covid positive and we identify a consequently large fraction of SARS-CoV-2-specific TCRs as HLA associated. Notably, this overlap provides probable pathogenic and HLA assignments to these TCRs, as demonstrated by the profound enrichment of these TCRs only among SARS-CoV-2-positive individuals expressing the appropriate restricting HLA. Thus, while only a subset of HLA-associated TCRs are observed in any given individual, the particular TCR subset observed reflects that individual’s history of exposure to many common antigens.

Our results imply that the vast majority of HLA-associated TCRs identified in this study likely derive from common antigens, making HLA-association a critical step in decoding the human T-cell repertoire. Moreover, the high imputation accuracy of our HLA models allows us to statistically HLA type all repertoires ever sequenced, thereby expanding the effective size of HLA-typed and TCR-sequenced cohorts by several orders of magnitude and further facilitating decoding efforts. The TCR repertoire is a Rosetta Stone of the human immune system, providing a rich source of information for characterizing both the genetic background and exposure history of individuals at a population scale. Mapping TCRs to HLAs and disease exposures and imputing HLA and disease exposure from TCRs represent important steps toward decoding the immunological history of individuals using immunosequencing.

## Materials & Methods

### Human Samples

TCR and HLA sequence data from human samples used for these studies were aggregated from several independent study collections described below. All necessary patient/participant consent has been obtained for each study and the appropriate institutional forms have been archived.

1. Whole blood samples from DLS (Discovery Life Sciences, Huntsville, AL) were collected under Protocol DLS13 for collection of clinical samples.
2. PBMC used for the sorted repertoire experiments were collected and processed by Bloodworks Northwest (Seattle, WA). Volunteer donors were consented and collected under the Bloodworks Research Donor Collection Protocol BT001.
3. PBMC were obtained from the Fred Hutchinson Cancer Research Center Research Cell Bank biorepository of healthy bone marrow donors. The sample collection protocol was approved and supervised by the Fred Hutchinson Cancer Research Center Institutional Review Board (IRB) (*33*).
4. Blood collected from human subjects were approved by the IRBs of Johns Hopkins University and Stanford University. All participants provided written informed consent prior to enrollment (*36*).
5. Blood collected for the ImmuneRACE Study has been approved by the Western IRB (WIRB) (reference number 1-1281891-1). The trial has received appropriate ethical approval from WIRB as described (*62*).
6. Procedures for the INCOV study were approved by the IRBs at Providence St. Joseph Health with IRB study number STUDY2020000175, the WIRB with IRB study number 20170658, and the University of Washington with IRB study numbers STUDY00000959 and STUDY00002929 (*63*).
7. Human samples from the Virology Research Clinic at the University of Washington was collected under an IRB-approved study (NCT04338360) (*45*).
8. Blood from the Institute of Medical, Pharmaceutical, and Health Sciences, Kanazawa University (KU) was collected with informed consent as described by documents approved by the KU review board (document number 585-2).

### Flow Cytometry and Cell Sorting

3*−*5*×*10^7^ PBMC were stained using a cocktail of antibodies that included CD3 (clone UCHT1, Biolegend), CD4 (clone OKT4, Biolegend), CD8 (clone RPA-T8, Biolegend), CD45RA(clone HI100, BD Bioscience), CCR7 (clone G043H7, Biolegend), CD95 (clone DX2, Biolegend), and CD28 (clone CD28.2, Biolegend) for 10 minutes at 4 degrees C. PBMC were washed with MACS buffer (Miltenyi Biotec) and then PE+ CD3+ T cells were enriched using anti-PE Microbeads with LS columns (Miltenyi Biotec) following the manufacturer’s protocol. The enriched CD3 sample was sorted using a FACSAria Fusion Flow Cytometer (BD Biosciences) to isolate naive CD4 (CD4^+^ CD3^+^ CD45RA^+^ CD95*^−^* CD28^+^ CCR7^+^), memory CD4 (CD4^+^ CD3^+^ non-naive CD4), naive CD8 (CD8^+^ CD3^+^ CD45RA^+^ CD95*^−^* CD28^+^ CCR7^+^) and memory CD8 (CD8^+^CD3^+^ non-naive CD8). Between 150, 000 and 3 *×* 10^6^ T cells were sorted and sent in for sequencing for each subset. TCR*β* sequencing was carried out using the ImmunoSEQ assay at Adaptive Biotechnologies.

### Model Training

We independently train a model for each HLA using the following procedure:

1. We randomly split 80% and 20% of all typed samples into a training and holdout set
2. Using typed HLA data, we label all samples as cases, controls or unlabeled for the HLA being modeled
3. We derive an ES set using the FET applied to cases and controls
4. We select a subset of ESs which associate to the HLA being modeled via the L1LR method
5. We derive an individual weight for each sequence in the list of ESs from step 3)
6. We generate two features which are the logarithm of the weighted sum of the convergent recombination count (i.e., number of unique DNA rearrangements mapping to a given CDR3) of enhanced sequences and the total unique rearrangements in the repertoire, respectively
7. We fit the standard Scikit-Learn logistic regression classifier to the training data and evaluate the model in cross validation to select the best FET p-value cutoff
8. We train the final model on all the train data and evaluate on the holdout set

Below we describe training steps in detail using the following definitions: Let *H* be the set of *N_H_* typed HLA allotypes in our training set and *h_i_ ∈ H*, *i* = 1*, …, N_H_* refer to any single HLA in the set. The training set of *h_i_* is denoted as *t_i_∈ T* where *T* is the set of *N_T_* training samples. We similarly define a test (or holdout) set for each HLA *t_i_^′^ ∈ T ^′^* where *T ^′^* is the set of *N_T_ ′* holdout samples. Below we drop the subscript *i* unless needed for clarity as the procedure is applied to each HLA independently.

**Step 1:** *T* and *T ^′^* are derived from a random 80/20 split of all typed data with *N_T_* + *N_T_ ′* = 4, 144. Fig. S1 shows the age, sex and ethnicity distribution of sample subjects.

**Step 2:** When building a model for a given HLA *h_i_*, samples are labeled as cases (label = 1) or controls (label = 0) if a subject expresses or does not express *h_i_*, respectively. A subset of controls are unlabeled if a p-group matched HLA to *h_i_*is expressed by the subject. P-group matched HLAs share the same antigen binding domain and thus may share T-cell receptor (TCR) specificities.

**Step 3:** For any HLA *h*, an initial set of ESs, *E*, is defined using Fisher’s Exact Test (FET) (*64*) applied to training set *t* of HLA *h*. The a range of p-value thresholds are used to identify ESs and the threshold is treated as a hyperparameter in the modeling; it is independently derived for each HLA we model.

**Step 4:** If HLA *h* is in linkage disequilibrium (LD) with other HLAs, the enhanced sequences specific to HLAs in LD will be present in the ES set of HLA *h* because of their co-occurrence in subjects (this is how LD is defined). The goal of our procedure is to identify and remove these sequences from the final set of ESs associated to *h*. Thus, a fundamental assumption of the model is that for a given train set, an ES may be associated with only a single HLA.

We enforce the assumption that each TCR is specific to one HLA by fitting a logistic regression model with L1 regularization and we refer to this as the L1LR method. We independently associate each ES in *E* with a single HLA. Thus, the assumption that a given TCR is associated to a single HLA is not globally enforced, i.e., it is possible that due to variations in training data that the same TCR may be associated to multiple HLAs though this is rare (see Fig. S2).

When associating individual ESs to HLAs, we model the presence/absence of any given ES *e_j_∈ E* in the training set *t* as a logistic regression classification. Here *j* denotes the individual TCRs in the ES set. For any sample *t_l_ ∈ t*, where *l* denotes individual samples in the training set *t*, the label for sequence *e_j_* is either 0 or 1 if the sequences is present or absent in *t_l_*, respectively. The feature vector for associating a given ES *e_j_*includes the indicator vector for the typed set of HLAs, i.e. *I_kl_*= 1 if sample *l* expresses HLA *h_k_* and *I_kl_* = 0 if sample *l* does not express HLA *h_k_*. Here, *k* indicates the subset of HLAs in *H* is in LD with HLA *h* (defined by FET p-value *<* 10*^−^*^3^). Thus, the feature vector is a binary vector indicating whether a subject has or does not have a given HLA which is observed to co-occur with the HLA being modeled. When associating sequences we include the total number of unique rearrangements in each sample repertoire, *d_l_* as a covariate not subject to L1 regularization. We model each ES independently and thus drop the subscript *j* in the following for clarity. Taken together a single ES *e* is modeled such that

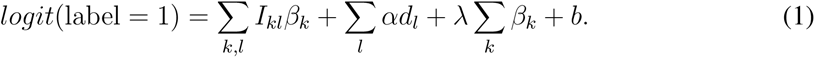

Here, *α* and *β* are free parameters, *b* is the bias term and *λ* is the regularization strength applied to *β*. We find the best-fit parameters by minimizing the log-loss and tune *λ* to be the smallest value that yields precisely one non-zero coefficient in *β_k_*. If the non-zero coefficient is *β_k_*_=_*_i_*, then the sequence is associated with the HLA being modeled and is retained as part of the final ES set. If the non-zero coefficient is *β_k̸_*_=_*_i_* the sequences is excluded from the final ES set of HLA *h* which is denoted as *Ẽ*. We weight each sample by the square root of the convergent recombination count when finding the best-fit parameters and zero count samples are given a weight of unity. The effectiveness of our procedure to resolve LD is demonstrated in Fig. S2.

**Step 5:** We derive a per-sequence weight, *w_j_* for each sequence *e_j_ ∈ Ẽ* by fitting a two feature logistic regression classifier. Here *j* refers to individual sequences in *Ẽ*. Each sequence is fit independently. For each sequence the target of the classifier is whether the sample is a case or control of HLA *h_i_* and the features are the convergent recombination count of the sequence and total number of unique rearrangements in the given repertoire sample. Essentially, we build a model to discriminate cases from controls using the count of a single sequence and take the coefficient as a weight. In detail, we standardize the features by subtracting the mean and normalizing to the standard deviation of the features across all samples. We fit the default logistic regression model in the Scikit-learn library which includes a fixed amount of L2 regularization (*λ* = 1). This procedure yields a weight for each sequence in *e_j_∈ Ẽ*.

**Step 6:** The final classifier model is a two feature model where the features *F* for subject *l* are

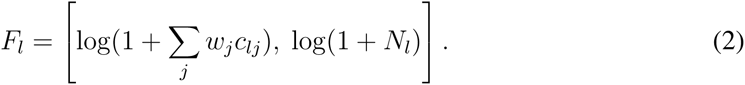

Here *c_lj_*the convergent recombination count of *e_j_* in sample *l*. These features, *F_l_*, are used in a the standard Scikit-learn logistic regression to predict the presence/absence of an HLA for a given sample.

**Step 7:** To model any given HLA, we fit the standard Scikit-Learn logistic regression classifier using features generated in Step 6). We treat the p-value cutoff for identifying ESs via the FET as described in Step 3) as a hyperparameter of each HLA model. We train and evaluate a model using a five-fold cross validation strategy for p-value cutoffs = [10*^−^*^3^, 10*^−^*^4^, 10*^−^*^5^, 10*^−^*^6^, 10*^−^*^7^, 10*^−^*^8^]. We adopt p-value cutoff of the highest performing cross-validation model.

**Step 8:** We generate our final model by adopting the p-value threshold determined in step 7) and train on the full set of training data. We evaluate the final model on the holdout set (Fig. S3-S8).

## Acknowledgments

This paper is dedicated to the memory of Peter Jacob Robert Ebert (1978–2023). He marched to his own drumbeat and was unconcerned with popular opinions. His intellectual curiosity, independence and rigor made him an innovative scientific leader and a wonderful collaborator. He was not only an incredibly clever scientist, but also a funny and kind colleague who will be missed by all who had the privilege to know him.

We thank Mary Carrington for discussion and helpful feedback which improved the manuscript and Thomas Snyder for guidance during early stages of manuscript drafting. We also thank the anonymous reviewers who provided helpful feedback that improved the clarity of the manuscript.

## Author Contributions

Conceptualization: HJZ, RT, PE, IC, JG, HR,JMC

Methodology: HJZ, CG, JL, LP, JG, JMC

Investigation: HJZ, RT, PE, IC, JG, JMC

Visualization: HJZ

Data Generation: MR, RE, HT

Supervision: JMC, HR, JG, WZ

Writing—original draft: HJZ, RT, PE, IC

Writing—review and editing: HJZ, RT, IC, JMC, JG, HR

## Data and Materials Availability

The derived data products are available in the Supplementary Materials in various tables. The sequence data are not publicly available as these data are part of a commercial product owned by Adaptive Biotechnologies.

## Footnotes

1 Throughout the manuscript, we refer to TCR*β* simply as TCR.

2 http://hla.alleles.org/alleles/pgroups.html

3 Degenerate subunit pairs are indicated by one of the degenerate subunits shown in parenthesis.

4 F1 score is the harmonic mean of precision and recall and useful metric of classification accuracy when there is large class imbalance such as the one we have for HLA imputation.

5 Errors represent 1 standard deviation and are bootstrapped.

6 We note that HLA-DR heterodimers are not subject to such pairing because the *α* chain is nearly invariable and thus these heterodimers behave similarly to class I HLAs.

7 These data do not have sequencing based HLA typing.

8 Our group has explored such analyses in disease contexts and found consistent results using TCR*α* or TCR*β* repertoires.

