## Supplementary Information for "Large-scale statistical mapping of T-cell receptor *β* sequences to Human Leukocyte Antigens"

### Supplementary Materials

#### S1. Degenerate HLAs

A subset of HLAs we model that are typed as distinct HLAs are specific to the same set of TCRs. Fig. 2A and 2B shows an example of such a degenerate HLA pair: A23:01 and A*23:17. Table 1 identifies degenerate HLAs.

#### S2. TCR Specificity and Three-Field Resolution

We explicitly test this assumption for class I HLAs by testing whether TCRs more strongly associate to the HLA group (one-field designation) rather than the two-field and whether synonymous substitutions impact TCR specificity. Class II HLAs are a two protein heterodimer which makes the analysis much more complicated.

We first aim to determine whether the set of TCRs associated with a given HLA depends on the three-field designation. Thus, we restrict our analysis to allotypes that have sufficient number of samples which differ in their three-field designation. A*68:01 and B*52:01 are the only class I HLAs with two well represented allotypes differing in their three-field designation. We identify ESs for each of these synonymously substituted allotypes (e.g., A*68:01:01 and A*68:01:02) independently and compare the fraction of ESs which overlap between the two sets. Because of statistical fluctuations resulting from sample-to-sample variation, we only expect partial overlap of the ES sets. Thus, we compare the overlap fraction for allotypes differing in their three-field designation to a null distribution derived via random partitioning of the same samples into two groups irrespective of their three-field designation. Fig. S9 demonstrates that the overlap fraction of TCRs is independent of the three-field designation confirming that a difference in the three-field designation does not appear impact TCR specificity for these HLAs. Given the third field designates synonymous substitutions, we expect this is a general result for all HLAs.

#### S3. Homozygosity across all loci

We test the hypothesis that homozgysity at one loci may yield elevated breadth at another class-matched loci due to competition across loci in surface expression and/or TCR binding. We do a pairwise comparison between all class-matched loci where subjects are either heterozygous or homozygous for one of the two loci. For example, we take all subjects that are heterozygous and homozygous for HLA-A and calculate the breadth observed in allotypes of HLA-B and vice versa. We do this for all six possible combinations of pairwise comparisons across the three class I and class II loci. We aggregate all pairwise comparisons and show the aggregated zscore distribution in Fig. S10.

We find the difference in the mean zscore for class I and class II HLAs is $0.10 \pm 0.03$ and $0.09 \pm 0.02$. While these differences are statistically signficant, they are an order of magnitude smaller than the increased breadth observed in homozygous subjects and are likely the result of correlated zygosity across class matched loci which we confirm via a Fisher's Exact Test. We note that correlation is especially strong between HLA-B and -C and HLA-DQ and -DR as expected given these loci are in particularly strong linkage disequilibrium. Despite correlation in zygosity, we find a very small difference and thus conclude that there does not appear to be competition across loci either for surface expression and/or TCR binding.

| Reported | Degenerate |
|---|---|
| A*23:01 | A*23:17 |
| DPA1*01:03+DPB1*03:01 | DPA1*01:03+DPB1*104:01 |
| DPA1*02:01+DPB1*17:01 | DPA1*02:01+DPB1*131:01 |
| DQA1*02:01+DQB1*02:02 | DQA1*02:01+DQB1*02:01 |
| DQA1*03:03+DQB1*03:01 | DQA1*03:01+DQB1*03:01 |
| DQA1*03:01+DQB1*03:02 | DQA1*03:03+DQB1*03:02 |
| DQA1*03:03+DQB1*02:02 | DQA1*03:03+DQB1*02:01 |
| DQA1*05:05+DQB1*03:01 | DQA1*05:01+DQB1*03:01, |
| DQA1*05:05+DQB1*03:01 | DQA1*05:05+DQB1*03:19 |
| DQA1*05:01+DQB1*02:01 | DQA1*05:05+DQB1*02:01 |
| DRB4*01:03 | DRB4*01:01 |
| DRB5*02:02 | DRB5*02:21 |

Table 1: **List of degenerate HLAs.** We are unable to distinguish between these sets of HLAs using TCR based typing as they have identical TCR specificities. When modeling degenerate HLAs, we only report the most commonly occurring HLA in the set.

| HLA | N Cases | Train AUCROC | Train CV AUCROC | Holdout AUCROC |
|---|---|---|---|---|
| A*01:01 | 763 | 0.99 | 0.99 | 1.00 |
| A*02:01 | 1314 | 1.00 | 1.00 | 1.00 |
| A*02:02 | 74 | 1.00 | 1.00 | 1.00 |
| A*02:05 | 75 | 1.00 | 1.00 | 0.99 |
| A*02:06 | 37 | 1.00 | 0.97 | 1.00 |
| A*03:01 | 703 | 1.00 | 1.00 | 0.99 |
| A*11:01 | 354 | 1.00 | 1.00 | 0.99 |
| A*23:01 | 231 | 1.00 | 1.00 | 1.00 |
| A*23:17 | 31 | 1.00 | 1.00 | 1.00 |
| A*24:02 | 495 | 1.00 | 0.99 | 0.99 |
| A*25:01 | 107 | 1.00 | 1.00 | 1.00 |

Continued on next page

| | | | | |
|---|---|---|---|---|
| A*26:01 | 144 | 1.00 | 1.00 | 0.99 |
| A*29:02 | 196 | 1.00 | 1.00 | 1.00 |
| A*30:01 | 174 | 1.00 | 1.00 | 1.00 |
| A*30:02 | 133 | 1.00 | 1.00 | 1.00 |
| A*31:01 | 177 | 1.00 | 1.00 | 1.00 |
| A*32:01 | 179 | 1.00 | 1.00 | 1.00 |
| A*33:01 | 74 | 1.00 | 1.00 | 0.99 |
| A*33:03 | 142 | 1.00 | 1.00 | 1.00 |
| A*34:02 | 60 | 1.00 | 1.00 | 0.99 |
| A*36:01 | 58 | 1.00 | 1.00 | 1.00 |
| A*66:01 | 40 | 0.99 | 0.98 | 1.00 |
| A*68:01 | 204 | 1.00 | 1.00 | 1.00 |
| A*68:02 | 160 | 1.00 | 1.00 | 1.00 |
| A*74:01 | 111 | 1.00 | 0.99 | 1.00 |
| B*07:02 | 660 | 1.00 | 1.00 | 1.00 |
| B*08:01 | 559 | 1.00 | 0.99 | 1.00 |
| B*13:02 | 125 | 1.00 | 1.00 | 1.00 |
| B*14:01 | 56 | 1.00 | 0.99 | 1.00 |
| B*14:02 | 175 | 1.00 | 1.00 | 1.00 |
| B*15:01 | 291 | 1.00 | 0.99 | 1.00 |
| B*15:03 | 104 | 1.00 | 1.00 | 1.00 |
| B*15:10 | 72 | 1.00 | 1.00 | 1.00 |
| B*15:16 | 36 | 1.00 | 1.00 | 1.00 |
| B*18:01 | 261 | 1.00 | 1.00 | 1.00 |
| B*27:05 | 169 | 1.00 | 1.00 | 0.99 |
| B*35:01 | 390 | 1.00 | 1.00 | 1.00 |
| B*35:02 | 53 | 1.00 | 0.98 | 0.96 |
| B*35:03 | 91 | 1.00 | 1.00 | 1.00 |
| B*37:01 | 65 | 1.00 | 1.00 | 1.00 |
| B*38:01 | 85 | 1.00 | 0.99 | 0.99 |
| B*39:01 | 71 | 1.00 | 1.00 | 1.00 |
| B*40:01 | 290 | 1.00 | 1.00 | 1.00 |
| B*40:02 | 80 | 1.00 | 1.00 | 0.99 |
| B*42:01 | 94 | 1.00 | 1.00 | 1.00 |
| B*44:02 | 405 | 1.00 | 1.00 | 0.99 |
| B*44:03 | 333 | 1.00 | 1.00 | 1.00 |
| B*45:01 | 114 | 1.00 | 1.00 | 1.00 |
| B*49:01 | 123 | 1.00 | 1.00 | 1.00 |
| B*50:01 | 78 | 1.00 | 0.99 | 1.00 |

Continued on next page

| | | | | |
|---|---|---|---|---|
| B*51:01 | 269 | 1.00 | 1.00 | 0.99 |
| B*52:01 | 74 | 1.00 | 1.00 | 1.00 |
| B*53:01 | 199 | 1.00 | 1.00 | 1.00 |
| B*55:01 | 95 | 1.00 | 1.00 | 1.00 |
| B*56:01 | 32 | 0.99 | 0.96 | 0.83 |
| B*57:01 | 166 | 1.00 | 0.99 | 0.99 |
| B*57:03 | 54 | 1.00 | 0.99 | 1.00 |
| B*58:01 | 109 | 1.00 | 1.00 | 1.00 |
| B*58:02 | 80 | 1.00 | 0.99 | 0.99 |
| C*01:02 | 208 | 0.99 | 0.95 | 0.96 |
| C*02:02 | 209 | 0.99 | 0.97 | 0.94 |
| C*02:10 | 119 | 1.00 | 0.98 | 0.97 |
| C*03:02 | 59 | 1.00 | 0.95 | 1.00 |
| C*03:03 | 282 | 0.96 | 0.89 | 0.89 |
| C*03:04 | 480 | 0.98 | 0.95 | 0.96 |
| C*04:01 | 829 | 0.99 | 0.99 | 0.98 |
| C*05:01 | 446 | 0.99 | 0.96 | 0.95 |
| C*06:02 | 559 | 1.00 | 1.00 | 0.99 |
| C*07:01 | 763 | 1.00 | 0.98 | 0.99 |
| C*07:02 | 757 | 1.00 | 0.96 | 0.97 |
| C*07:04 | 73 | 1.00 | 0.66 | 0.68 |
| C*07:18 | 81 | 0.98 | 0.96 | 0.93 |
| C*08:01 | 30 | 1.00 | 0.95 | 0.87 |
| C*08:02 | 246 | 1.00 | 0.99 | 0.99 |
| C*12:02 | 55 | 1.00 | 0.92 | 0.94 |
| C*12:03 | 246 | 1.00 | 1.00 | 0.98 |
| C*14:02 | 75 | 1.00 | 0.98 | 0.99 |
| C*15:02 | 114 | 0.99 | 0.84 | 0.93 |
| C*15:05 | 43 | 0.99 | 0.93 | 0.98 |
| C*16:01 | 320 | 1.00 | 0.99 | 0.98 |
| C*17:01 | 132 | 1.00 | 0.94 | 0.94 |

Table 2: **Model performance of class I HLAs.** The HLA allele and number of subjects with the given HLA are listed in columns 1 and 2, respectively. Columns 3-5 indicate model performance when the trained model is evaluated on the train, train CV and holdout data sets, respectively.

| HLA | N Cases | Train AUCROC | Train CV AUCROC | Holdout AUCROC | In LD |
|---|---|---|---|---|---|
| DPA1*01:03+DPB1*01:01 | 387 | 1.00 | 0.99 | 0.99 | No |
| DPA1*01:03+DPB1*02:01 | 796 | 1.00 | 0.99 | 1.00 | Yes |
| DPA1*01:03+DPB1*02:02 | 39 | 1.00 | 0.97 | 0.97 | Yes |
| DPA1*01:03+DPB1*03:01 | 415 | 0.99 | 0.99 | 0.99 | Yes |
| DPA1*01:03+DPB1*04:01 | 1618 | 1.00 | 0.99 | 0.99 | Yes |
| DPA1*01:03+DPB1*04:02 | 571 | 1.00 | 1.00 | 1.00 | Yes |
| DPA1*01:03+DPB1*05:01 | 117 | 1.00 | 1.00 | 0.99 | No |
| DPA1*01:03+DPB1*06:01 | 85 | 1.00 | 0.99 | 1.00 | Yes |
| DPA1*01:03+DPB1*104:01 | 101 | 1.00 | 1.00 | 1.00 | Yes |
| DPA1*01:03+DPB1*105:01 | 66 | 0.99 | 0.93 | 1.00 | No |
| DPA1*01:03+DPB1*10:01 | 70 | 1.00 | 0.98 | 0.95 | No |
| DPA1*01:03+DPB1*11:01 | 92 | 1.00 | 0.96 | 0.99 | No |
| DPA1*01:03+DPB1*13:01 | 85 | 1.00 | 1.00 | 1.00 | No |
| DPA1*01:03+DPB1*14:01 | 55 | 1.00 | 0.99 | 1.00 | No |
| DPA1*01:03+DPB1*17:01 | 78 | 1.00 | 0.97 | 0.99 | No |
| DPA1*01:03+DPB1*18:01 | 113 | 1.00 | 1.00 | 1.00 | Yes |
| DPA1*02:01+DPB1*01:01 | 461 | 1.00 | 0.99 | 0.99 | Yes |
| DPA1*02:01+DPB1*02:01 | 172 | 0.92 | 0.57 | 0.58 | No |
| DPA1*02:01+DPB1*03:01 | 71 | 1.00 | 0.99 | 0.95 | No |
| DPA1*02:01+DPB1*04:01 | 324 | 1.00 | 0.57 | 0.61 | No |
| DPA1*02:01+DPB1*04:02 | 96 | 1.00 | 0.54 | 0.57 | No |
| DPA1*02:01+DPB1*05:01 | 36 | 1.00 | 0.94 | 1.00 | No |
| DPA1*02:01+DPB1*09:01 | 39 | 0.99 | 0.94 | 0.99 | Yes |
| DPA1*02:01+DPB1*105:01 | 52 | 1.00 | 0.80 | 0.79 | No |
| DPA1*02:01+DPB1*10:01 | 84 | 1.00 | 0.99 | 0.98 | Yes |
| DPA1*02:01+DPB1*11:01 | 145 | 1.00 | 0.99 | 1.00 | Yes |
| DPA1*02:01+DPB1*131:01 | 39 | 1.00 | 1.00 | 1.00 | Yes |
| DPA1*02:01+DPB1*13:01 | 139 | 1.00 | 1.00 | 1.00 | Yes |
| DPA1*02:01+DPB1*14:01 | 71 | 1.00 | 1.00 | 1.00 | Yes |
| DPA1*02:01+DPB1*17:01 | 143 | 1.00 | 1.00 | 1.00 | Yes |
| DPA1*02:01+DPB1*18:01 | 35 | 1.00 | 0.70 | 0.73 | No |
| DPA1*02:02+DPB1*01:01 | 268 | 1.00 | 1.00 | 0.99 | Yes |
| DPA1*02:02+DPB1*02:01 | 63 | 1.00 | 0.61 | 0.56 | No |
| DPA1*02:02+DPB1*04:01 | 78 | 0.97 | 0.80 | 0.84 | No |
| DPA1*02:02+DPB1*05:01 | 102 | 1.00 | 1.00 | 0.95 | Yes |
| DPA1*02:06+DPB1*05:01 | 34 | 1.00 | 0.96 | 1.00 | Yes |
| DPA1*03:01+DPB1*01:01 | 54 | 1.00 | 0.78 | 0.91 | Yes |
| DPA1*03:01+DPB1*105:01 | 140 | 1.00 | 1.00 | 1.00 | Yes |

Continued on next page

| | | | | | |
|---|---|---|---|---|---|
| DQA1*01:01+DQB1*02:01 | 65 | 1.00 | 0.80 | 0.83 | No |
| DQA1*01:01+DQB1*02:02 | 78 | 1.00 | 0.81 | 0.72 | No |
| DQA1*01:01+DQB1*03:01 | 122 | 1.00 | 0.78 | 0.75 | No |
| DQA1*01:01+DQB1*03:02 | 53 | 1.00 | 0.76 | 0.68 | No |
| DQA1*01:01+DQB1*05:01 | 602 | 1.00 | 0.99 | 1.00 | Yes |
| DQA1*01:01+DQB1*06:02 | 81 | 1.00 | 0.91 | 0.98 | No |
| DQA1*01:01+DQB1*06:03 | 32 | 1.00 | 0.94 | 0.98 | No |
| DQA1*01:02+DQB1*02:01 | 161 | 1.00 | 0.81 | 0.85 | No |
| DQA1*01:02+DQB1*02:02 | 181 | 1.00 | 0.80 | 0.77 | No |
| DQA1*01:02+DQB1*03:01 | 214 | 1.00 | 0.72 | 0.73 | No |
| DQA1*01:02+DQB1*03:02 | 102 | 1.00 | 0.69 | 0.76 | No |
| DQA1*01:02+DQB1*03:03 | 41 | 0.99 | 0.54 | 0.55 | No |
| DQA1*01:02+DQB1*03:19 | 59 | 1.00 | 0.86 | 0.91 | No |
| DQA1*01:02+DQB1*04:02 | 63 | 1.00 | 0.81 | 0.65 | No |
| DQA1*01:02+DQB1*05:01 | 221 | 1.00 | 0.99 | 0.99 | No |
| DQA1*01:02+DQB1*05:02 | 149 | 1.00 | 1.00 | 1.00 | Yes |
| DQA1*01:02+DQB1*05:03 | 33 | 1.00 | 0.85 | 0.90 | No |
| DQA1*01:02+DQB1*06:02 | 830 | 1.00 | 1.00 | 1.00 | Yes |
| DQA1*01:02+DQB1*06:03 | 70 | 1.00 | 0.97 | 0.99 | No |
| DQA1*01:02+DQB1*06:04 | 184 | 1.00 | 0.99 | 1.00 | Yes |
| DQA1*01:02+DQB1*06:09 | 135 | 1.00 | 1.00 | 1.00 | Yes |
| DQA1*01:03+DQB1*02:01 | 46 | 1.00 | 0.62 | 0.76 | No |
| DQA1*01:03+DQB1*02:02 | 43 | 1.00 | 0.64 | 0.59 | No |
| DQA1*01:03+DQB1*03:01 | 62 | 1.00 | 0.68 | 0.69 | No |
| DQA1*01:03+DQB1*03:02 | 33 | 1.00 | 0.62 | 0.56 | No |
| DQA1*01:03+DQB1*05:01 | 55 | 0.99 | 0.92 | 0.91 | No |
| DQA1*01:03+DQB1*06:01 | 54 | 1.00 | 1.00 | 1.00 | Yes |
| DQA1*01:03+DQB1*06:02 | 53 | 1.00 | 1.00 | 0.95 | No |
| DQA1*01:03+DQB1*06:03 | 299 | 1.00 | 1.00 | 0.99 | Yes |
| DQA1*01:04+DQB1*05:03 | 122 | 1.00 | 1.00 | 1.00 | Yes |
| DQA1*01:05+DQB1*05:01 | 145 | 1.00 | 1.00 | 1.00 | Yes |
| DQA1*01:05+DQB1*06:02 | 30 | 1.00 | 0.81 | 0.74 | No |
| DQA1*02:01+DQB1*02:01 | 74 | 1.00 | 1.00 | 1.00 | No |
| DQA1*02:01+DQB1*02:02 | 602 | 1.00 | 1.00 | 1.00 | Yes |
| DQA1*02:01+DQB1*03:01 | 131 | 1.00 | 0.99 | 0.98 | No |
| DQA1*02:01+DQB1*03:02 | 64 | 1.00 | 0.97 | 0.97 | No |
| DQA1*02:01+DQB1*03:03 | 164 | 1.00 | 0.99 | 0.98 | Yes |
| DQA1*02:01+DQB1*04:02 | 32 | 1.00 | 0.87 | 0.98 | No |
| DQA1*02:01+DQB1*05:01 | 109 | 1.00 | 0.77 | 0.66 | No |

Continued on next page

| | | | | | |
|---|---|---|---|---|---|
| DQA1*02:01+DQB1*06:02 | 110 | 1.00 | 0.96 | 0.97 | No |
| DQA1*02:01+DQB1*06:03 | 39 | 0.90 | 0.75 | 0.79 | No |
| DQA1*02:01+DQB1*06:04 | 30 | 1.00 | 0.78 | 0.93 | No |
| DQA1*03:01+DQB1*02:01 | 59 | 1.00 | 0.73 | 0.77 | No |
| DQA1*03:01+DQB1*02:02 | 49 | 1.00 | 0.97 | 0.99 | No |
| DQA1*03:01+DQB1*03:01 | 78 | 1.00 | 0.99 | 1.00 | No |
| DQA1*03:01+DQB1*03:02 | 442 | 1.00 | 0.99 | 1.00 | Yes |
| DQA1*03:01+DQB1*05:01 | 55 | 1.00 | 0.64 | 0.68 | No |
| DQA1*03:01+DQB1*06:02 | 55 | 1.00 | 0.81 | 0.68 | No |
| DQA1*03:02+DQB1*03:03 | 76 | 1.00 | 1.00 | 1.00 | Yes |
| DQA1*03:03+DQB1*02:01 | 52 | 0.98 | 0.90 | 0.99 | No |
| DQA1*03:03+DQB1*02:02 | 120 | 1.00 | 1.00 | 1.00 | Yes |
| DQA1*03:03+DQB1*03:01 | 306 | 1.00 | 1.00 | 1.00 | Yes |
| DQA1*03:03+DQB1*03:02 | 98 | 1.00 | 1.00 | 1.00 | Yes |
| DQA1*03:03+DQB1*03:03 | 40 | 1.00 | 0.92 | 0.82 | Yes |
| DQA1*03:03+DQB1*04:02 | 30 | 1.00 | 0.73 | 0.71 | No |
| DQA1*03:03+DQB1*05:01 | 72 | 1.00 | 0.73 | 0.81 | No |
| DQA1*03:03+DQB1*06:02 | 65 | 1.00 | 0.77 | 0.92 | No |
| DQA1*04:01+DQB1*02:02 | 38 | 1.00 | 0.72 | 0.79 | No |
| DQA1*04:01+DQB1*03:01 | 33 | 1.00 | 0.90 | 0.81 | No |
| DQA1*04:01+DQB1*03:19 | 64 | 1.00 | 1.00 | 1.00 | Yes |
| DQA1*04:01+DQB1*04:02 | 241 | 1.00 | 1.00 | 1.00 | Yes |
| DQA1*04:01+DQB1*05:01 | 38 | 1.00 | 0.66 | 0.57 | No |
| DQA1*04:01+DQB1*06:02 | 49 | 1.00 | 0.91 | 0.95 | No |
| DQA1*05:01+DQB1*02:01 | 639 | 1.00 | 1.00 | 1.00 | Yes |
| DQA1*05:01+DQB1*02:02 | 57 | 1.00 | 0.99 | 1.00 | No |
| DQA1*05:01+DQB1*03:01 | 122 | 1.00 | 1.00 | 1.00 | No |
| DQA1*05:01+DQB1*03:02 | 63 | 0.97 | 0.97 | 0.99 | No |
| DQA1*05:01+DQB1*03:03 | 30 | 1.00 | 0.93 | 0.95 | No |
| DQA1*05:01+DQB1*04:02 | 31 | 1.00 | 0.69 | 0.90 | No |
| DQA1*05:01+DQB1*05:01 | 80 | 1.00 | 0.73 | 0.72 | No |
| DQA1*05:01+DQB1*06:02 | 95 | 1.00 | 0.83 | 0.88 | No |
| DQA1*05:01+DQB1*06:03 | 37 | 1.00 | 0.70 | 0.80 | No |
| DQA1*05:05+DQB1*02:01 | 84 | 1.00 | 1.00 | 1.00 | No |
| DQA1*05:05+DQB1*02:02 | 69 | 1.00 | 0.97 | 1.00 | No |
| DQA1*05:05+DQB1*03:01 | 601 | 1.00 | 1.00 | 0.99 | Yes |
| DQA1*05:05+DQB1*03:02 | 66 | 0.99 | 0.94 | 0.94 | No |
| DQA1*05:05+DQB1*03:19 | 116 | 1.00 | 1.00 | 1.00 | Yes |
| DQA1*05:05+DQB1*05:01 | 101 | 1.00 | 0.76 | 0.67 | No |

Continued on next page

| | | | | | |
|---|---|---|---|---|---|
| DQA1*05:05+DQB1*06:02 | 117 | 1.00 | 0.84 | 0.82 | No |
| DQA1*05:05+DQB1*06:03 | 41 | 1.00 | 0.73 | 0.77 | No |
| DQA1*06:01+DQB1*03:01 | 40 | 1.00 | 1.00 | 1.00 | Yes |
| DRB1*01:01 | 442 | 1.00 | 0.99 | 1.00 | Yes |
| DRB1*01:02 | 138 | 1.00 | 1.00 | 1.00 | Yes |
| DRB1*01:03 | 49 | 1.00 | 1.00 | 1.00 | Yes |
| DRB1*03:01 | 655 | 1.00 | 1.00 | 1.00 | Yes |
| DRB1*03:02 | 103 | 1.00 | 1.00 | 1.00 | Yes |
| DRB1*04:01 | 421 | 1.00 | 1.00 | 1.00 | Yes |
| DRB1*04:02 | 46 | 1.00 | 0.99 | 1.00 | Yes |
| DRB1*04:03 | 49 | 1.00 | 1.00 | 1.00 | Yes |
| DRB1*04:04 | 190 | 1.00 | 1.00 | 1.00 | Yes |
| DRB1*04:05 | 76 | 1.00 | 1.00 | 1.00 | Yes |
| DRB1*04:07 | 72 | 1.00 | 1.00 | 1.00 | Yes |
| DRB1*07:01 | 750 | 1.00 | 1.00 | 1.00 | Yes |
| DRB1*08:01 | 109 | 1.00 | 1.00 | 1.00 | Yes |
| DRB1*08:02 | 34 | 1.00 | 1.00 | 1.00 | Yes |
| DRB1*08:04 | 93 | 1.00 | 1.00 | 1.00 | Yes |
| DRB1*09:01 | 126 | 1.00 | 1.00 | 1.00 | Yes |
| DRB1*10:01 | 79 | 1.00 | 1.00 | 1.00 | Yes |
| DRB1*11:01 | 388 | 1.00 | 1.00 | 1.00 | Yes |
| DRB1*11:02 | 85 | 1.00 | 1.00 | 1.00 | Yes |
| DRB1*11:04 | 141 | 1.00 | 1.00 | 0.99 | Yes |
| DRB1*12:01 | 154 | 1.00 | 1.00 | 0.99 | Yes |
| DRB1*13:01 | 364 | 1.00 | 1.00 | 0.99 | Yes |
| DRB1*13:02 | 353 | 1.00 | 1.00 | 1.00 | Yes |
| DRB1*13:03 | 95 | 1.00 | 0.99 | 1.00 | Yes |
| DRB1*14:54 | 129 | 1.00 | 1.00 | 1.00 | Yes |
| DRB1*15:01 | 611 | 1.00 | 1.00 | 1.00 | Yes |
| DRB1*15:02 | 75 | 1.00 | 1.00 | 1.00 | Yes |
| DRB1*15:03 | 202 | 1.00 | 1.00 | 1.00 | Yes |
| DRB1*16:01 | 64 | 1.00 | 1.00 | 1.00 | Yes |
| DRB1*16:02 | 54 | 1.00 | 1.00 | 1.00 | Yes |
| DRB3*01:01 | 654 | 1.00 | 0.99 | 0.99 | Yes |
| DRB3*01:62 | 91 | 1.00 | 0.99 | 1.00 | Yes |
| DRB3*02:02 | 1005 | 1.00 | 1.00 | 1.00 | Yes |
| DRB3*03:01 | 374 | 1.00 | 1.00 | 1.00 | Yes |
| DRB4*01:01 | 361 | 1.00 | 1.00 | 1.00 | Yes |
| DRB4*01:03 | 1014 | 0.99 | 0.98 | 0.98 | Yes |

Continued on next page

| | | | | | |
|---|---|---|---|---|---|
| DRB5*01:01 | 677 | 1.00 | 1.00 | 1.00 | Yes |
| DRB5*01:02 | 41 | 1.00 | 1.00 | 1.00 | Yes |
| DRB5*02:02 | 75 | 1.00 | 1.00 | 1.00 | Yes |

Table 3: **Model performance of class II HLAs.** The HLA allele and number of subjects with the given HLA are listed in columns 1 and 2, respectively. Columns 3-5 indicate model performance when the trained model is evaluated on the train, train CV and holdout data sets, respectively. Column 6 indicates whether the heterodimer is observed to be in linkage disequilibrium. HLAs in linkage equilibrium likely result from trans-complementation of the subunits (see main text).

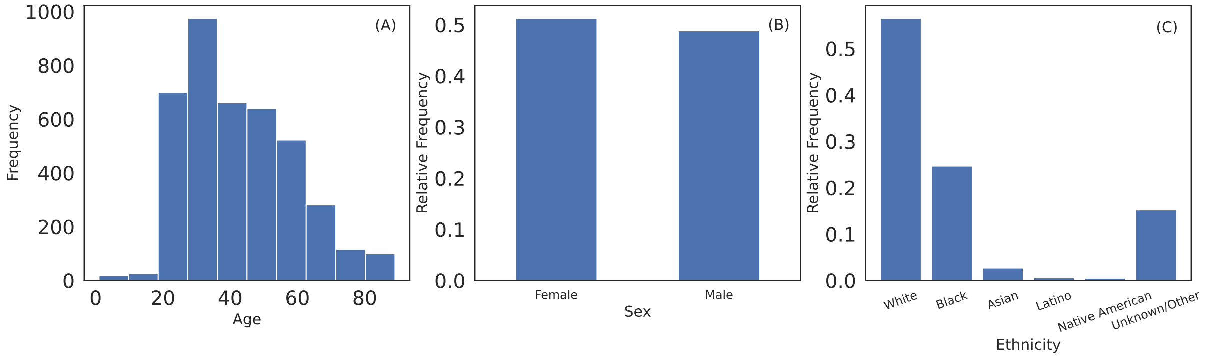


Fig S1: **Demographic distribution of sample population.** (A) Age, (B) sex and (C) self-reported ethnicity distribution of sample subjects.

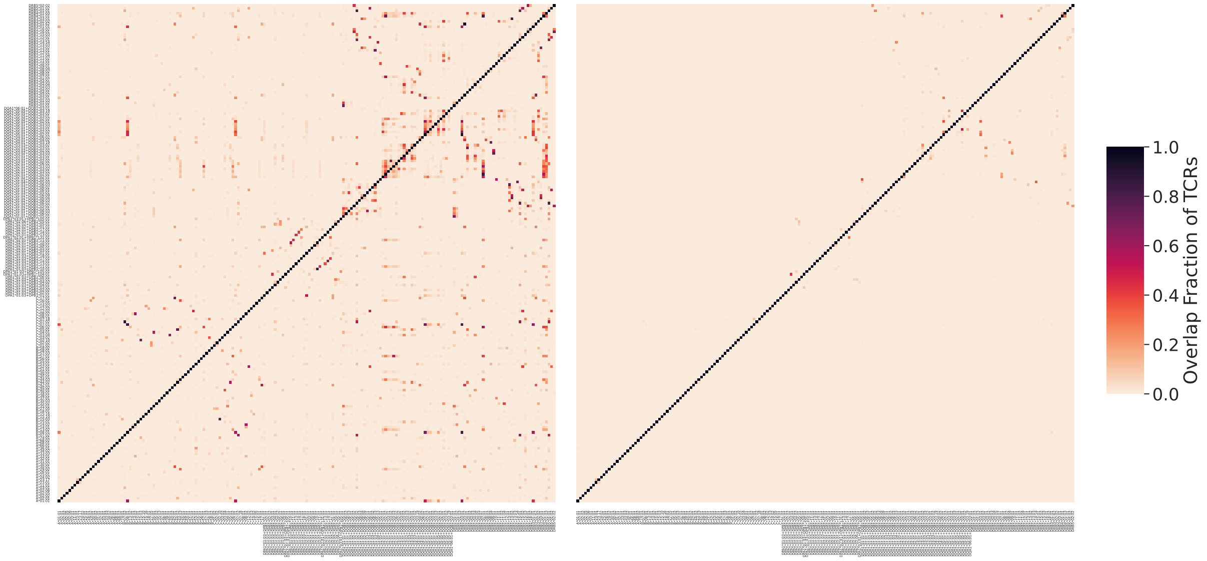


Fig S2: **Effectiveness of L1LR method to resolve LD.** Heatmap showing the amount of TCR sharing before (left) and after (right) applying L1LR method to resolve LD. The color indicates the fractional sharing of TCRs amongst pairs of HLAs relative to the total number of TCRs associated with the HLA indicated on the vertical axis. After accounting for LD, we observe significantly less sharing of TCRs amongst HLA allotypes. The remaining shared TCRs may be the result of LD that is too strong to resolve using our method or true sharing of TCRs between multiple HLAs (e.g., as shown in the main text).

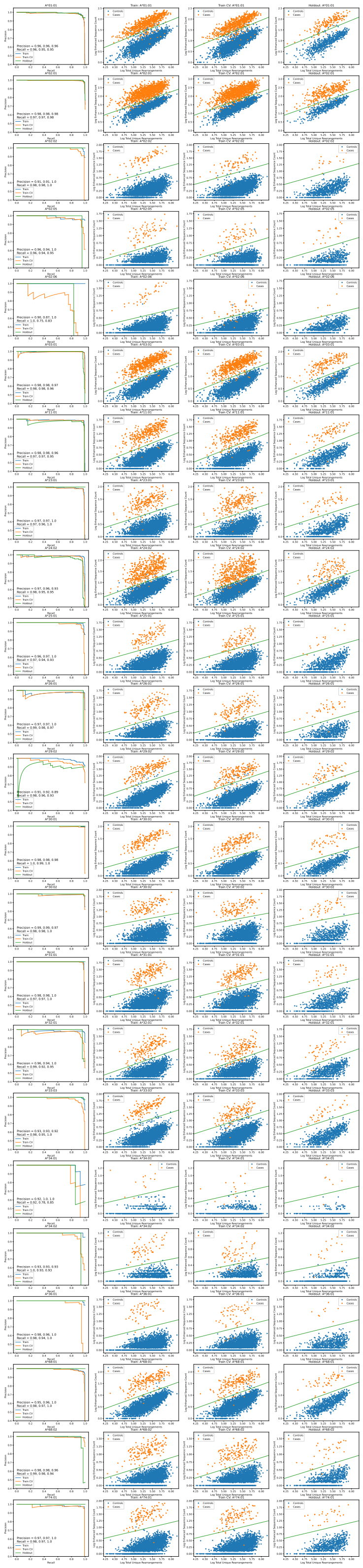

Fig S3: **HLA-A models.** Left column shows the precision-recall curve for the trained model evaluated on the train sample (middle left), train sample in cross-validation (middle right) and on the holdout sample (right) for HLA-A. The green line indicates the calibration threshold.

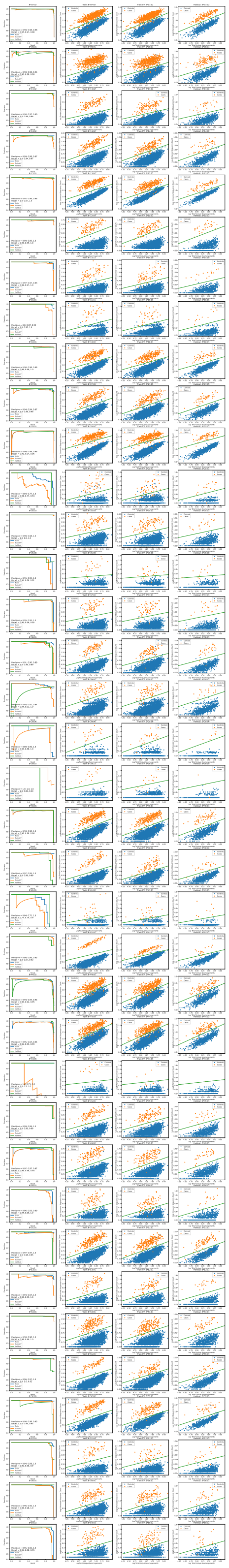

Fig S4: **HLA-B models.** Left column shows the precision-recall curve for the trained model evaluated on the train sample (middle left), train sample in cross-validation (middle right) and on the holdout sample (right) for HLA-B. The green line indicates the calibration threshold.

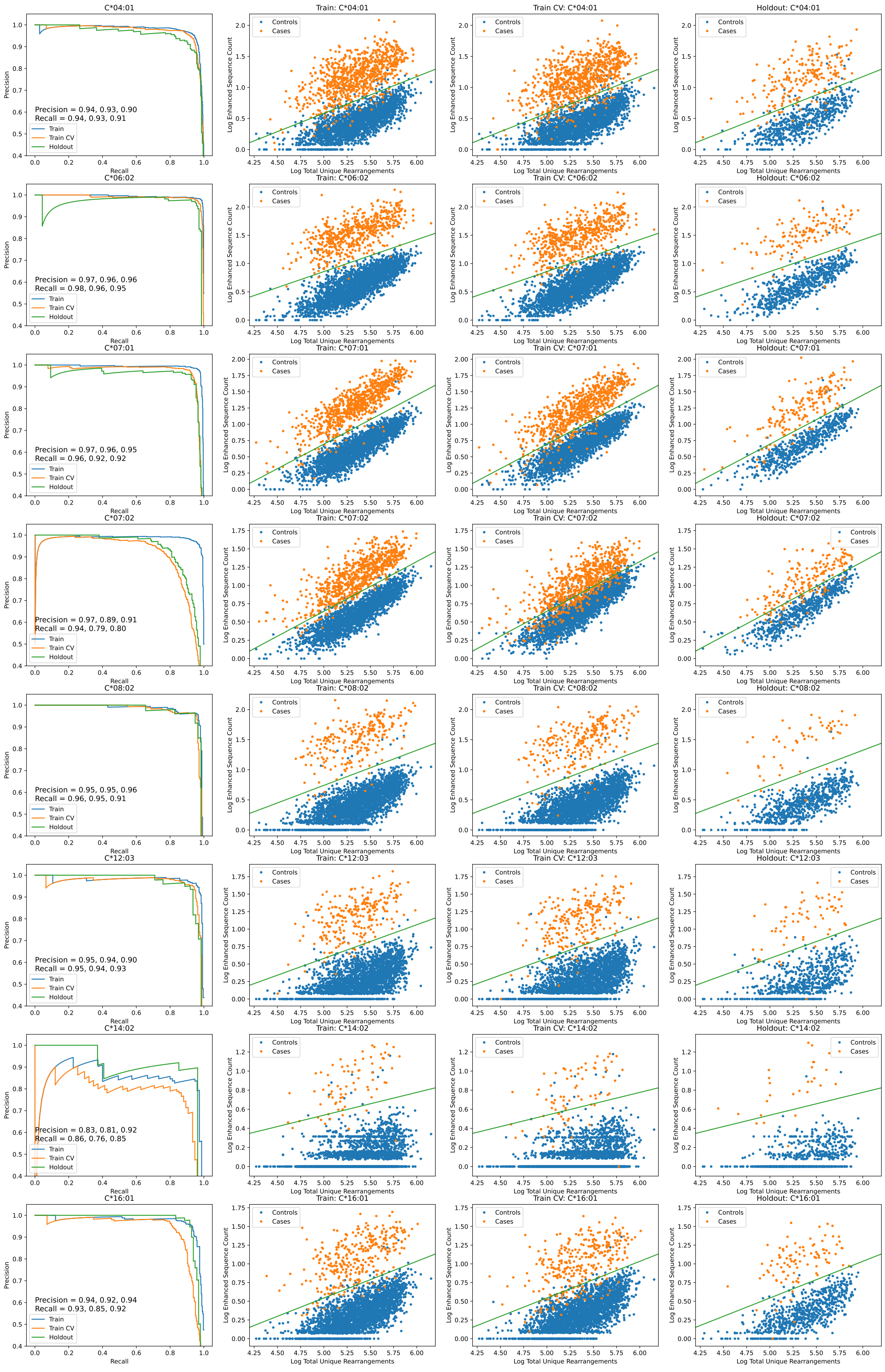


Fig S5: **HLA-C models.** Left column shows the precision-recall curve for the trained model evaluated on the train sample (middle left), train sample in cross-validation (middle right) and on the holdout sample (right) for HLA-C. The green line indicates the calibration threshold.

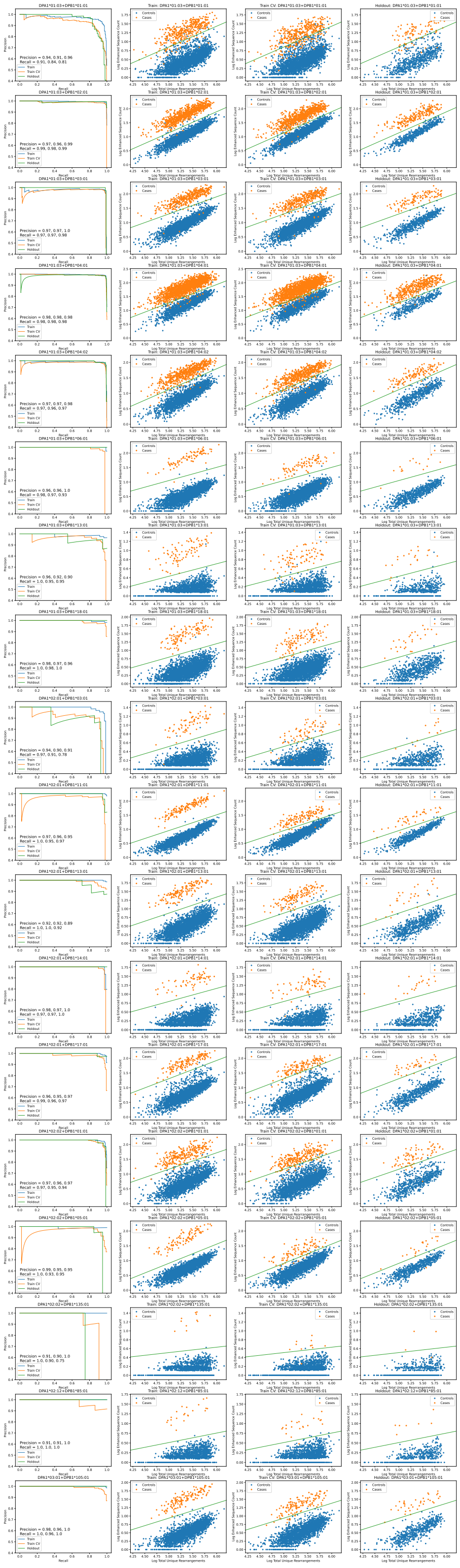

Fig S6: **HLA-DP models.** Left column shows the precision-recall curve for the trained model evaluated on the train sample (middle left), train sample in cross-validation (middle right) and on the holdout sample (right) for HLA-DP. The green line indicates the calibration threshold.

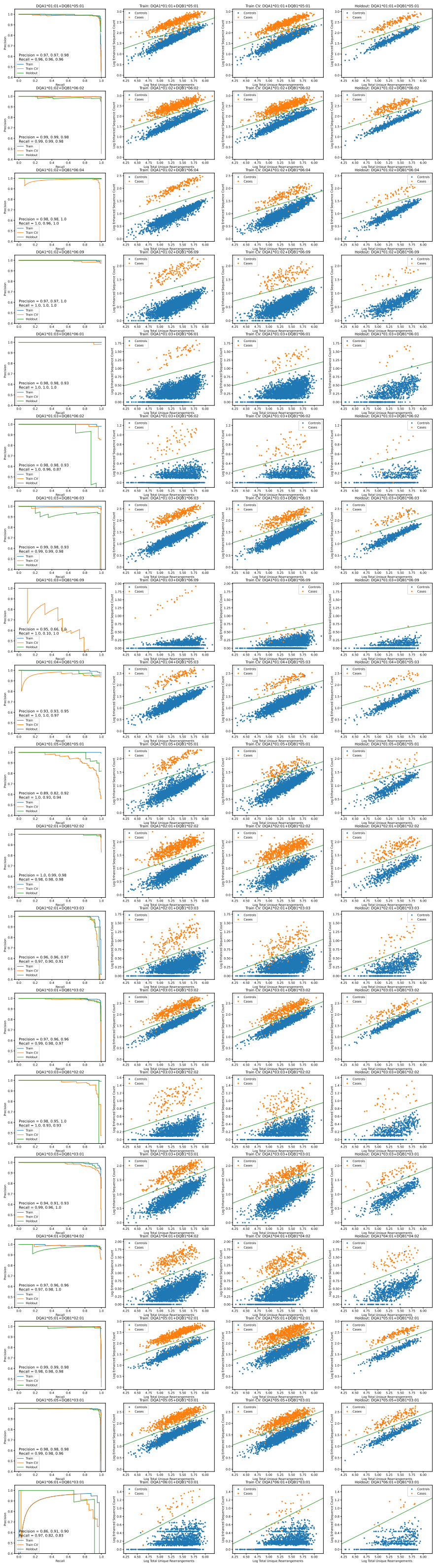

Fig S7: **HLA-DQ models.** Left column shows the precision-recall curve for the trained model evaluated on the train sample (middle left), train sample in cross-validation (middle right) and on the holdout sample (right) for HLA-DQ. The green line indicates the calibration threshold.

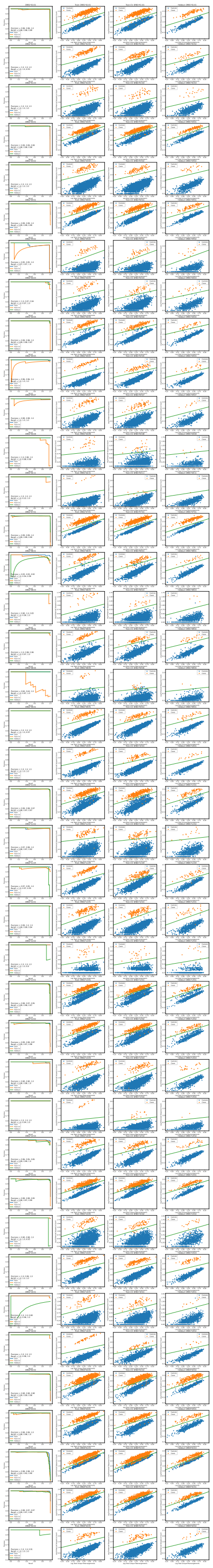

Fig S8: **HLA-DR models.** Left column shows the precision-recall curve for the trained model evaluated on the train sample (middle left), train sample in cross-validation (middle right) and on the holdout sample (right) for HLA-DR. The green line indicates the calibration threshold.

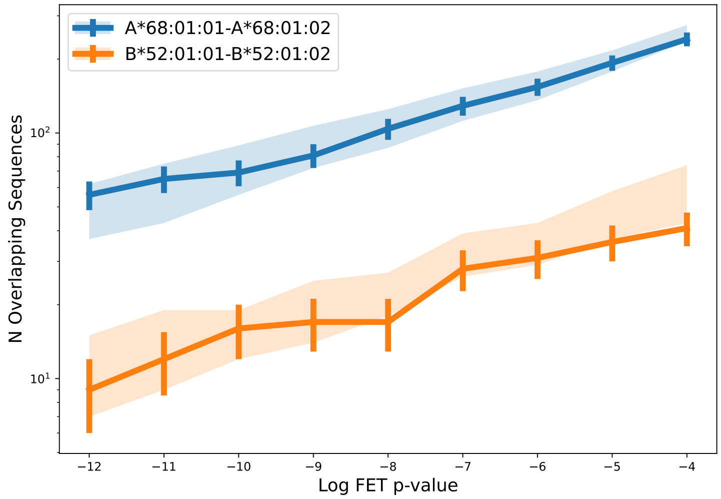


Fig S9: **TCRs specificity apparently independent of synonymous mutations in coding regions of HLAs.** Number of TCRs overlapping in two ES sets of HLAs differing only in their third field designation as a function of the FET p-value threshold used to identify ESs. The solid line shows the overlap in two sets of ESs identified using the three field designation to divide samples into two groups. Error bars are poisson uncertainties. Even if the TCR specificity is identical for HLAs differing only in their third field designation, we expect differences in the ES sets due to statistical fluctuations resulting from sampling. To account for this effect, we compare the observed overlap fraction when segregating samples by third field designation to a null distribution of overlap fractions derived from 100 realizations of two samples which are randomly divided irrespective of the third field designation. The two randomly divided samples in each realization are selected such that the relative size of the two samples matches the size of two samples split by the three field designation. Shaded regions indicate the distribution of the overlap fraction from these 100 realizations. Consistency between the overlap fraction of ESs derived from samples differing in their three field resolution and ones based on random permutation demonstrates that TCR specificity is independent of three field resolution for the two HLA allotypes tested.

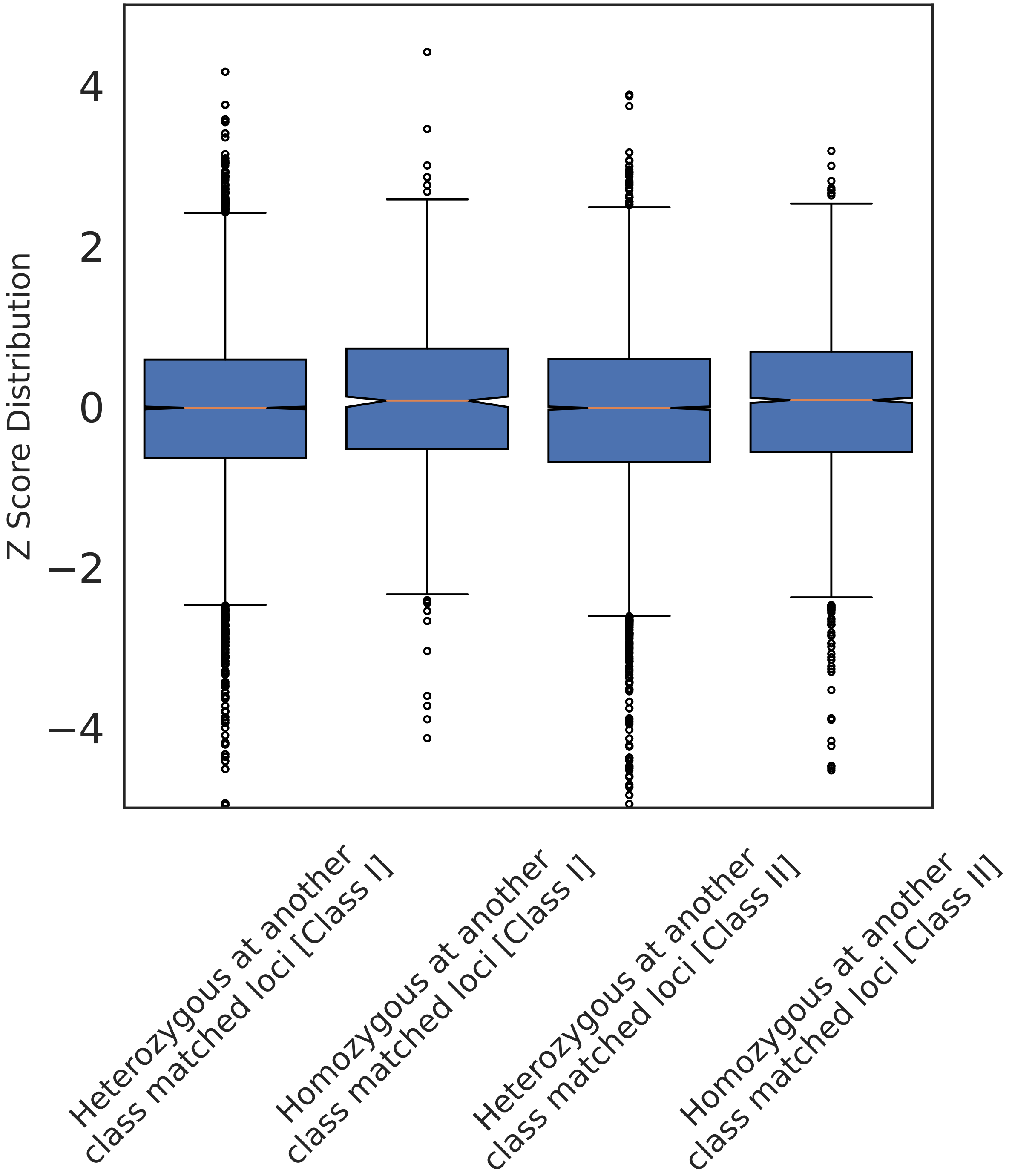


Fig S10: **We test the hypothesis that homozygosity at one loci increases breadth at another class matched loci.** The distribution of zscores across class matched loci from a pairwise loci comparison. We calculate the zscore of the breadth at one loci in the case where another class matched loci is heterozygous or homozygous. We aggregate all pairwise comparisons for class I and class II separately.
